# High levels of frataxin overexpression leads to mitochondrial and cardiac toxicity in mouse models

**DOI:** 10.1101/2020.03.31.015255

**Authors:** Brahim Belbellaa, Laurence Reutenauer, Nadia Messaddeq, Laurent Monassier, Hélène Puccio

## Abstract

Friedreich ataxia (FA) is currently an incurable inherited mitochondrial disease caused by reduced levels of frataxin (FXN). Cardiac dysfunction is the main cause of premature death in FA. AAV-mediated gene therapy constitutes a promising approach for FA, as demonstrated in cardiac and neurological mouse models. While the minimal therapeutic level of FXN protein to be restored and biodistribution have recently been defined for the heart, it is unclear if FXN overexpression could be harmful. Indeed, depending on the vector delivery route and dose administrated, the resulting FXN protein level could reach very high levels in the heart, cerebellum, or in off-target organs such as the liver. The present study demonstrates safety of FXN cardiac overexpression up to 9-fold the normal endogenous level, but significant toxicity to the mitochondria and heart above 20-fold. We show gradual severity with increasing FXN overexpression, ranging from subclinical cardiotoxicity to left ventricle dysfunction. This appears to be driven by impairment of mitochondria respiratory chain, ultrastructure and homeostasis, which lead to myofilaments alteration, cell death and fibrosis. Overall, this study underlines the need, during the development of gene therapy approaches, to consider appropriately vector potency, long term safety and biomarkers to monitor such events.

## INTRODUCTION

Friedreich ataxia (FA) is a rare neurodegenerative disease characterized by spinocerebellar and sensory ataxia associated with hypertrophic cardiomyopathy (HCM) ^1^. Cardiac dysfunction is the main cause of premature death in FA patients ^2^. Nighty-five percent of FA patient present a homozygous (GAA) expansion within the first intron of the frataxin gene (*FXN*)^3^. This pathological expansion causes the heterochromatinization of the locus leading to reduced transcription of the *FXN* gene ^4^. FXN is a highly conserved mitochondrial protein regulating the biosynthesis of iron sulfur clusters (Fe-S) through an interaction with the *de novo* ISC complex assembly machinery ^5^. Fe-S are prosthetic groups crucial for many biological functions, including the mitochondrial respiratory chain and iron metabolism ^6^. Frataxin deficiency leads to dysregulation of Fe-S biogenesis, impairment of Fe-S enzymes, mitochondrial dysfunction, iron metabolism dysregulation and eventually to cellular dysfunction and death ^7, 8^.

The therapeutic potential of AAV-mediated *in vivo* gene therapy in preventing and rescuing mitochondrial dysfunction associated with FXN deficiency in both cardiac and neurological tissues has clearly been demonstrated in several mouse studies ^9-11^. To translate successfully these initial proof-of-concept studies to the clinic, we undertook to define the therapeutic thresholds conditioning the rescue of the cardiac phenotype, as well as the potential toxic effects associated with FXN overexpression. Recently, using the *Mck* mouse, a conditional knockout (cKO) model that recapitulates most features of the FA cardiomyopathy ^7^, we demonstrated that the therapeutic outcome of AAV-FXN gene therapy is directly and proportionally correlated with the vector biodistribution in the heart and that the correction of only half the cardiomyocyte was sufficient to restore fully the cardiac morphology and function ^12^. In contrast to earlier reports in transgenic mouse models overexpression FXN ^13, 14^, several recent studies conducted *in vitro* ^15, 16^ and with transgenic drosophila ^17, 18^ have suggested that constitutive FXN overexpression could be deleterious.

In the present study, we investigated whether *in vivo* AAV-mediated gene transfer could be detrimental in wild-type mice or frataxin-deficient *Mck* mice. We evaluated FXN overexpression *in vivo* and its consequences on mitochondria and heart function and morphology. FXN protein expression over 20-fold the endogenous level is toxic for the mitochondria and results in severe impairment of complexes I and II enzymatic activity, alteration of the mitochondria ultrastructure, cardiomyocytes cell death and fibrosis leading to heart dysfunction. While the mitochondrial unfolded protein response did not appear to be activated, the integrated stress response (ISR) was strongly induced, with the overexpression of FGF21 and Galectin-3 genes encoding for plasma-secreted proteins. Overexpression of frataxin up to 9-fold the endogeneous level in the cardiac tissue was uneventful, suggesting that the maximum safe level of FXN overexpression is between 9 and 20-fold the normal level. Strikingly, liver overexpression of FXN >90-fold the normal level did not result in any detectable toxicity, suggesting organs specific susceptibility to FXN overexpression.

## RESULTS

### FXN overexpression leads to impairment of mitochondria SDH activity

Endogenous FXN protein ([mFXN]) levels in the heart ventricles of wild-type (WT) animals was 147±42 ng/mg. In the previous dose-response study^12^, the concentration of human frataxin protein ([hFXN-HA]), assayed by ELISA, ranged from 2 to 10,927 ng per mg of total heart protein, after intravenous administration of AAVRh.10-CAG-hFXN-HA vector with doses from 5×10^13^ down to 1×10^12^ vg/kg. Interestingly, heart tissue sections with very high level of hFXN overexpression systematically showed colocalization of SDH enzymatic activity impairment at hotspots of frataxin expression (Figure S1). This was particularly obvious in the animal displaying the highest-level of vector copies (7.68 per diploid genome on average (VCN)) and protein concentration (10,927 ng/mg or 74-fold the endogenous level). No SDH impairment was observed in the cardiomyocytes expressing much lower hFXN-HA levels or neighboring these hotspots, nor was it observed in *Mck* mice treated with lower vector doses expressing much lower levels of hFXN-HA. Interestingly, the cardiomyocytes with high hFXN expression and impaired SDH activity did not show any iron accumulation following Perls-DAB staining, as is typically seen in FXN deficiency and/or Fe-S biosynthesis impairment (Figure S1)^7, 19^. Furthermore, these hotspots were not associated with cardiomyocyte cell death, fibrosis or cell-infiltrations, as assessed by hematoxylin & eosin (H&E) and wheat germ agglutinin (WGA) staining of adjacent heart tissue sections (Figure S1). Importantly, the left ventricle function of these mice did not appear impaired at rest. Their left ventricle (LV) shortening fraction (SF) and cardiac blood output normalized to body weight (CO/BW) were corrected to WT levels (Figure S2A-B), similarly to *Mck* mice treated with a 2-fold lower vector dose. Most likely, this is explained by the relatively small proportion of cardiomyocytes and heart surface affected overall (<10-20%). Indeed, we showed previously that the functional rescue of the cardiac phenotype is achieved with as low as 50-60% of heart surface corrected ^12^. It is worth noting that these mice did not display significant LV hypertrophy (Figure S2C), but the LV diameters at the end systole (LV-ESD) and diastole (LV-EDD) appeared slightly higher in the mouse treated with 5×10^13^vg/kg and expressing 74-fold the normal level of FXN (Figure S2D-E). Overall, these preliminary results suggested that very high level of hFXN-HA protein in cardiomyocytes could lead to impaired mitochondrial SDH enzymatic activity but without notable iron accumulation.

### Overexpression of hFXN in WT mice leads to mitochondrial and cardiac toxicity

In order to rule out any potential confounding effects from the *Mck* mouse cardiac phenotype, we undertook to replicate in WT mice these experimental conditions, i.e. VCN ≥ 7 and [hFXN] ≥ 10,000 ng/mg (Figure 1A-C). Therefore, 7-weeks old WT mice received 5×10^14^vg/kg or 5×10^13^vg/kg of AAVRh.10-CAG-hXN-HA vector. Compared to age-matched *Mck* mice, one-log higher vector dose was required in WT mice to achieve the same vector biodistribution and overexpression. However, the vector potency was similar in both mouse models, when the [hFXN] was normalized to VCN (Figure S3). At 21 weeks of age, i.e. 3 months post-injection, all treated WT mice were alive, with normal body weight growth curve (Figure S4A) and no behavioral sign of stress or suffering. Upon sacrifice, no gross anatomical anomalies were observed at the levels of the heart, lung, liver, kidney, digestive tract, or skeletal muscles.

**Figure 1.**
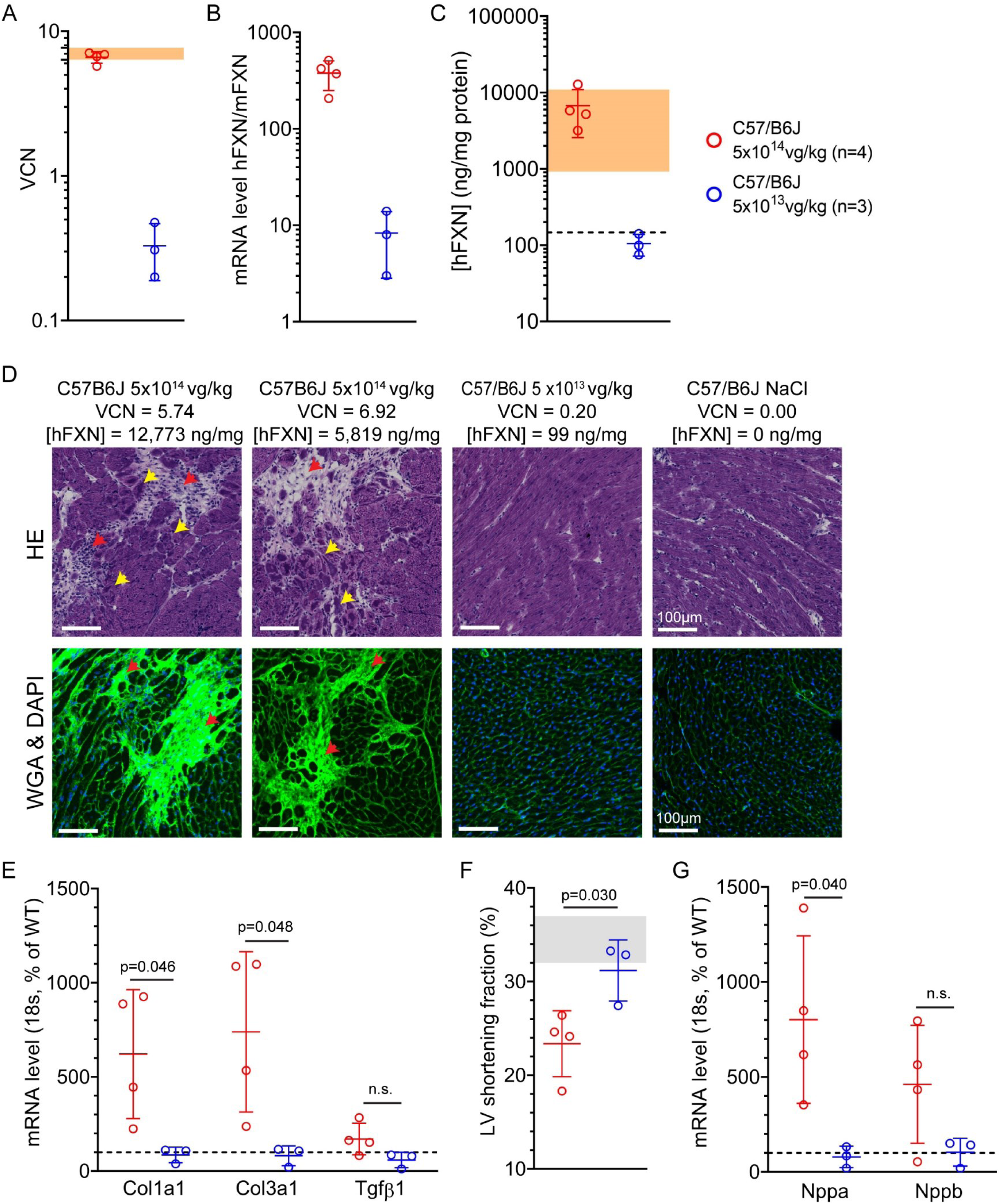
High level of FXN overexpression in the heart of wild-type mice treated with AAVRh.10-CAG-hFXN-HA vector is associated with cardiac fibrosis and subclinical impairment of heart function and morphology. Wild-type (WT) C57/B6J mice were treated at 7-weeks of age with vehicle or AAVRh.10-CAG-hFXN-HA vector, at the dose of 5×10^14^ (n=4, red) or 5×10^13^ (n=3, blue) vg/kg and sacrificed at 21 weeks. **(A)** qPCR quantification of vector copies per diploid genome (VCN) in heart tissue. Light orange area represents VCN range observed previously in *Mck* mice and illustrated in Figure S1. **(B)** RTqPCR quantification of hFXN mRNA level normalized to mouse FXN (mFXN) mRNA level. **(C)** ELISA assay quantification of human FXN protein concentration ([hFXN]) in the heart, normalized to mg of total heart protein. Light orange area represents [hFXN] range observed previously in *Mck* mice and illustrated in Figure S1. Blacked-dotted line represent normal [mFXN] endogenous level in WT mice (147±42 ng/mg). **(D)** Histological analysis of heart fibrosis and cell infiltrates by hematoxylin and eosin (HE) or wheat-germ-agglutinin (WGA) staining. Representative image of the LV anterior wall. Red arrows indicate fibrosis and cell infiltrates and yellow arrows cardiomyocytes displaying subcellular disorganization. VCN and [hFXN] values are indicated above respectively for each animal. Scale bar, 100µm. **(E)** RTqPCR quantification of Col1a1, Col3a1 and Tgfβ1 mRNA levels in the heart, normalized to 18S and reported as percentage of WT level. **(F)** Echocardiography assessment of left ventricle (LV) shortening fraction at 21 weeks of age. Grey area represents the range of normal values in control mice. Grey-shaded area corresponds to normal values range. **(G)** RTqPCR quantification of Nppa and Nppb mRNA levels in the heart, normalized to 18S and reported as percentage of WT. Blacked-dotted line represents normal mRNA level in control mice. Data are represented as mean ± SD. Student t-test and p values are reported, with n.s. p>0.05.

The consequences of hFXN-HA overexpression on the heart histology were evaluated following H&E and WGA staining of adjacent heart frozen tissue sections (Figures 1D and S5). Both staining revealed sparse fibrotic patches in the three animals treated with 5×10^14^vg/kg and expressing the highest levels of hFXN-HA, between 5,206 and 12,773 ng of FXN per mg of protein (i.e. 34 and 85-fold the endogenous protein level in the heart). The heart fibrosis, quantified as the accumulation of extracellular matrix, covered between 1.92 and 3.96% of the heart tissue section surface in these mice, while animals with lower hFXN-HA levels or injected with vehicle had values ranging from 1.49 to 2.05% (Figure S6A). This overlap between the two dose groups reflects the variability of vector biodistribution and expression, especially in the high dose group. These observations were supported by the increased genes expression of Col1a1, Col3a1 and Tgfβ1, indicative of ongoing fibrosis, more pronounced in the high dose group (Figure 1E). Overall, these results support the progressive gradation of the cardiotoxicity with the level of FXN overexpression.

While none of the mice developed severe cardiac dysfunction, the three animals expressing the highest hFXN-HA levels displayed meaningful reduction of LV shortening fraction (Figures 1F and S4B-C and Video S1) and heart hypertrophy (Figure S4E-F), but without decrease of the CO/BW at rest (Figure S4D). No functional nor morphologic heart alterations were observed in the mice treated with vehicle or with vector expressing hFXN-HA levels close to the endogenous level. These observations were line with the correlation analysis between the level of [hFXN-HA] and the cardiac dysfunction (Figure S4G-H) and with gene expression levels of *Nppa* and *Nppb*, biomarkers indicative of pressure/volume overload (Figure 1G).

To assess the mitochondria function, semiquantitative histoenzymatic assays were performed to probe the activity of the respiratory chain complexes I, II and IV, on adjacent heart tissue sections (Figures 2A, S5 and S7). The enzymatic activity of SDH was sparsely impaired (<10% of the heart surface) in the same three mice expressing the highest hFXN-HA levels but was unaltered in all other animals (Figures 2A, S5 and S6B). As observed initially in *Mck* mice (Figure S1), impaired SDH activity strictly co-localized with hotspots of hFXN-HA expression. Furthermore, the enzymatic activity of complex I (NADH) was also impaired sparsely (Figures 2A and S7) while the enzymatic activity of complex IV (COX) was not affected (Figures 2A and S5). Importantly, untreated *Mck* mice heart deficient for FXN also display impaired SDH and NADH enzymatic activities, two Fe-S cofactors dependent proteins, but not for COX which enzymatic activity is dependent on heme cofactors ^20^.

**Figure 2.**
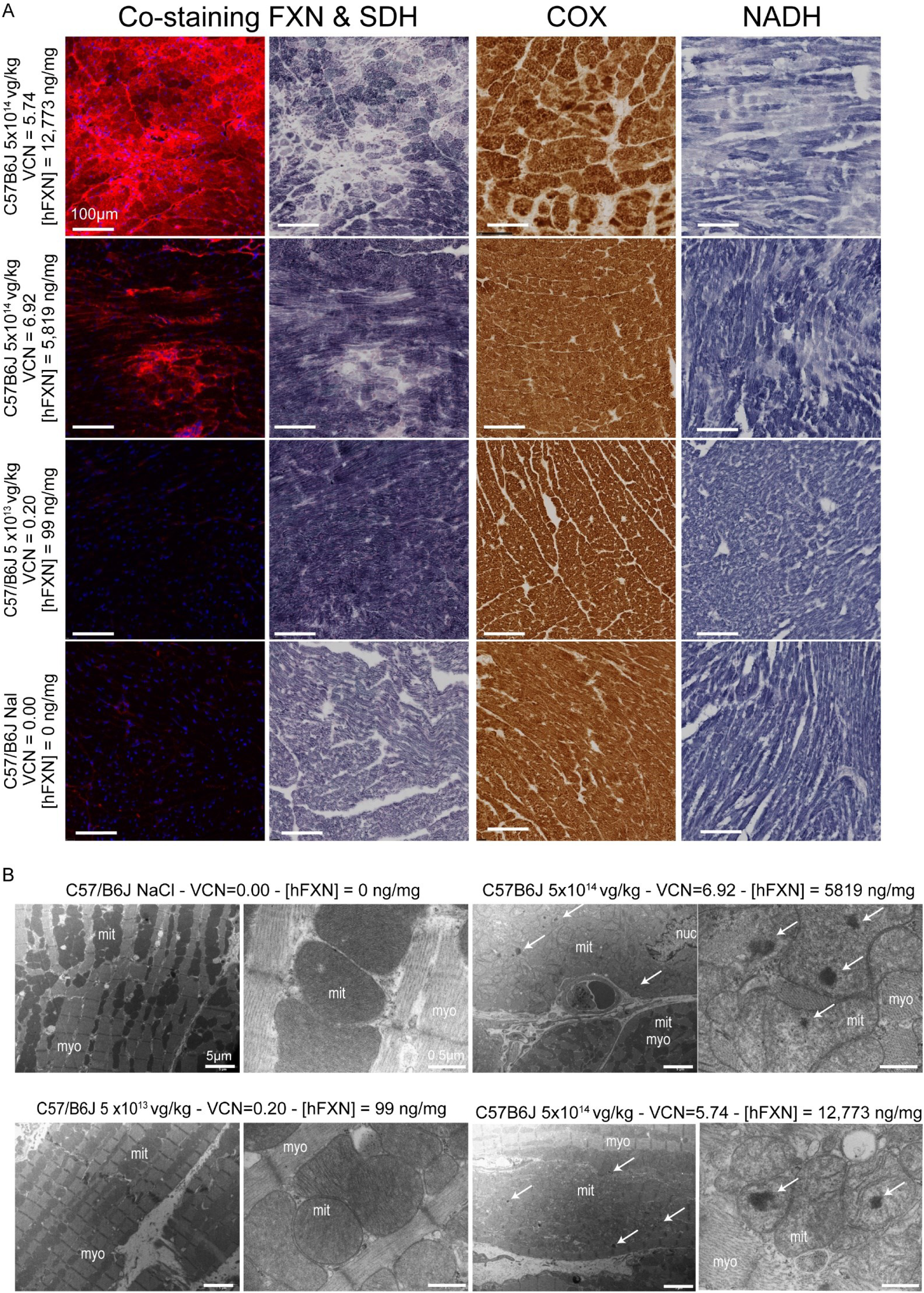
Impairment of mitochondrial ultrastructure and activity of respiratory chain complexes I and II following high level of FXN overexpression in the heart of wild-type mice treated with AAVRh.10-CAG-hFXN-HA vector. **(A)** Adjacent heart tissue sections from treated and control WT mice were collected to perform histological analyses to assess mitochondria respiratory chain complexes function I, II and IV. The two left columns represent the same tissue section and microscopy field co-stained for hFXN by immunofluorescent labelling (identical time exposure) and for succinate dehydrogenase (SDH) activity by histoenzymatic assay. The middle-right column represents cytochrome c oxidase (COX) histoenzymatic activity. The right-column represents NADH-ubiquinone oxidoreductase (NADH) histoenzymatic activity. Scale bars, 100µm. **(B)** Transmission electron micrographs of heart to assess cardiomyocytes and mitochondria ultrastructure. The same animals were analyzed in panel A and B. Representative images of the left ventricle myocardium acquired at low and high magnifications. White arrows indicate mitochondrial electron dense bodies; myo, myofibrils; mito, mitochondria; nuc, nucleus. The dose of vector administrated, VCN and [hFXN] are reported for each animal next to images series. Scale bars: low magnification 5µm, high magnification 0.5µm.

The consequences of FXN overexpression on the cardiomyocytes ultrastructure was assessed by transmission electron microscopy (TEM) (Figure 2B). In mice expressing the highest levels of hFXN-HA, a substantial proportion of cardiomyocytes displayed severe alterations in their subcellular organization, with scattered and disordered myofibrils. The mitochondria were swollen with few cristae whose stacking is crucial for mitochondria bioenergetic efficiency ^21^. We also frequently observed the presence of electron dense bodies inside the mitochondria matrix, not reminiscent of collapsed cristae nor iron deposits, as commonly observed in FA patients and untreated *Mck* mice cardiomyocytes ^7, 22, 23^. To rule-out the presence of mitochondrial iron deposits, adjacent ultrathin sections were stained with bismuth subnitrate to label iron stored in mitochondrial and cytosolic ferritins ^22, 24-26^. All mice displayed iron labelling in lysosome, as expected since this organelle is central to ferritin turn-over, but none showed mitochondrial iron deposits nor ferritin accumulation, independently of hFXN-HA levels or the presence of severely altered cardiomyocytes and mitochondria (Figures 2B and S8). Collectively, these observations rule out iron accumulation as part of the cardio- and mitochondrial toxicity mechanism, which is in line with the lack of iron deposits observed in treated *Mck* mice (Figure S1).

To further investigate the consequence of hFXN-HA overexpression on mitochondria homeostasis, autophagy, mitochondria biomass and mitochondrial bioenergetics were evaluated on adjacent heart tissue sections from the same WT mice. Analysis of prohibitin (Phb) and Sqstm1, respectively indicative of mitochondria biomass and cellular autophagy, revealed a substantial increase of both markers, specifically in cardiomyocytes where SDH enzymatic activity was impaired (Figure 3A-B). Similarly, the increase of mitochondrial proteins pan-lysine acetylation was specifically observed in SDH negative cardiomyocytes (Figure 3C). This post-translational modification is associated with the dysregulation of the NAD+/NADH and acetyl-CoA metabolisms ^27-29^, and results in the modulation of enzymatic activity and decreased bioenergetic efficiency in the mitochondria ^30, 31^. Strikingly, the neighboring cardiomyocytes, which are SDH positive, displayed normal levels for Phb, Sqstm1 and pan-lysine acetylation. This strongly suggests that the metabolic consequences of toxic FXN overexpression are mediated in a cell-autonomous manner. Furthermore, this was associated with the induction of the Nrf2-mediated oxidative stress response (Figure S9A) and the integrated stress response (ISR) (Figure S9B-C). Interestingly, the expression of ISR target genes encoding for secreted proteins (Fgf21 and Lgasl3) were also significantly elevated (Figure S9 D-G). These cytokines have been identified as biomarkers of mitochondria and heart dysfunction, in previous (pre)clinical studies ^32, 33^.

**Figure 3.**
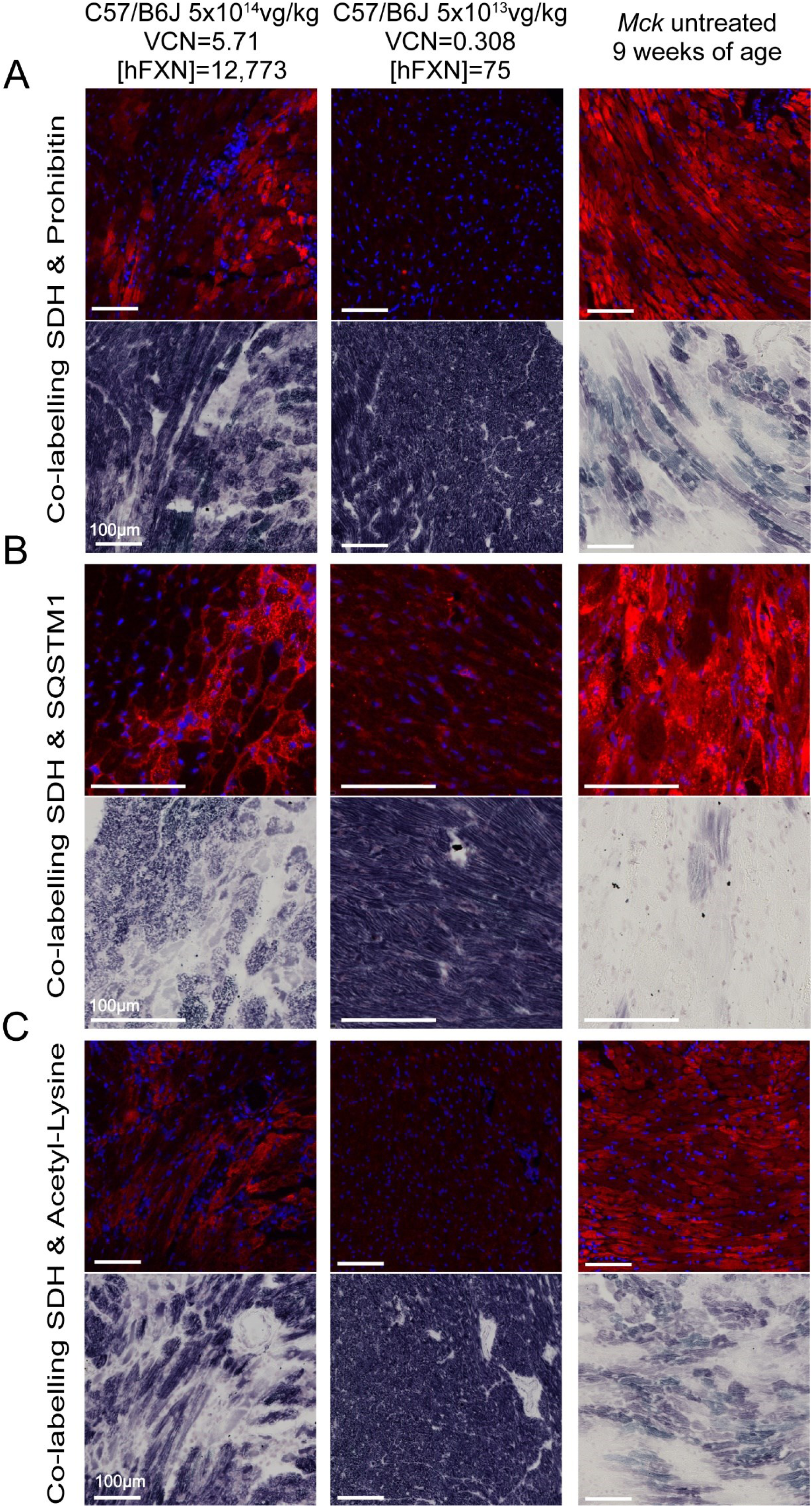
Toxic overexpression of FXN in the heart of WT mice is associated with increased mitochondrial proliferation, autophagy, and dysregulation of NADH/NAD+ metabolism. Heart tissue sections were co-labelled for biomarkers of mitochondrial homeostasis and for SDH enzymatic activity, as proxy for hFXN overexpression and mitochondrial function impairment. The same microscopy field was acquired for both labelling, to perform co-localization analysis. Representative images from the LV anterior wall are reported for WT C57/B6J mice treated with AAVRh.10-CAG-hFXN-HA vector at the dose of 5×10^14^ or 5×10^13^ vg/kg. The respective dose, VCN and [hFXN] are reported for each animal above each image series. Untreated *Mck* mice sacrificed at 9 weeks of age were used as positive controls of mitochondrial impairment and biomarkers elevation. **(A)** Co-staining of SDH enzymatic activity (lower row) and prohibitin (upper row, identical time exposure) which is indicative mitochondria biomass. **(B)** Co-staining of SDH enzymatic activity (lower row) and SQSTM1 (upper row, identical time exposure) which is indicative autophagy and mitophagy. **(C)** Co-staining of SDH enzymatic activity (lower row) and pan-lysine acetylation (upper row, identical time exposure) which is indicative NAD+/NADH or bioenergetic imbalance. DNA labeled with DAPI. Scale bar, 100μm.

Altogether, these results suggest that cardiac overexpression of hFXN-HA ≥20-30-fold the normal endogenous level is toxic to the heart and mitochondria. The mitochondria toxicity is characterized by severe alterations of the mitochondria ultrastructure, bioenergetics, homeostasis and the induction of mitochondria stress responses.

### High level of FXN overexpression results in acute cardiotoxicity and compromised the gene therapy outcome

While the study in WT mice recapitulated and confirmed the mitochondrial toxicity observed initially in *Mck* mice, it was unclear if this toxicity could be related in part to the high dose of vector administrated, the resulting high number of vector copies per cell, and/or the HA tag fused in C-terminal of the transgene (Figure S10A).

To address these questions, we designed an optimized expression cassette, AAVRh.10-hFXN (Figure S10B), with increased potency (Figure S3) and without the HA tag. When administered to *Mck* mice, AAVRh.10-hFXN vector resulted in much higher levels of hFXN expression in heart and in liver despite similar VCN values (Figures 4A-B and S10C-E). Interestingly, the initial vector had similar potency in the heart of WT and *Mck* mice (Figure S3), despite the significant transcriptional and metabolic dysregulations occurring upon FXN depletion ^7, 9, 12, 34^. In contrast, the optimized vector showed in average a 34-fold higher potency in *Mck* mice than in WT mice (Figure S3). This increased potency is likely driven by the strong induction of the ISR and Eif2α phosphorylation in *Mck* mice heart^34^, which might favorably impact the transcription and translation of AAVRh.10-hFXN vector, in part through the optimized Kozak sequence. In line with this hypothesis, both vectors displayed similar transcription efficiency in the liver of *Mck* mice (Figure S11A-B), which is not depleted in FXN and does not present any functional or molecular phenotype. Nonetheless, the optimization of the Kozak sequence still leads by itself in a 48-fold increase of the average hFXN protein levels in the liver (Figure S11C-D). Furthermore, the optimized vector displayed much higher potency and hFXN protein levels in the *Mck* mice heart than liver, when quantified by ELISA assay (Figures 4B and S11C) or compared side-by-side using Western-Blot analysis (Figure S11D).

**Figure 4.**
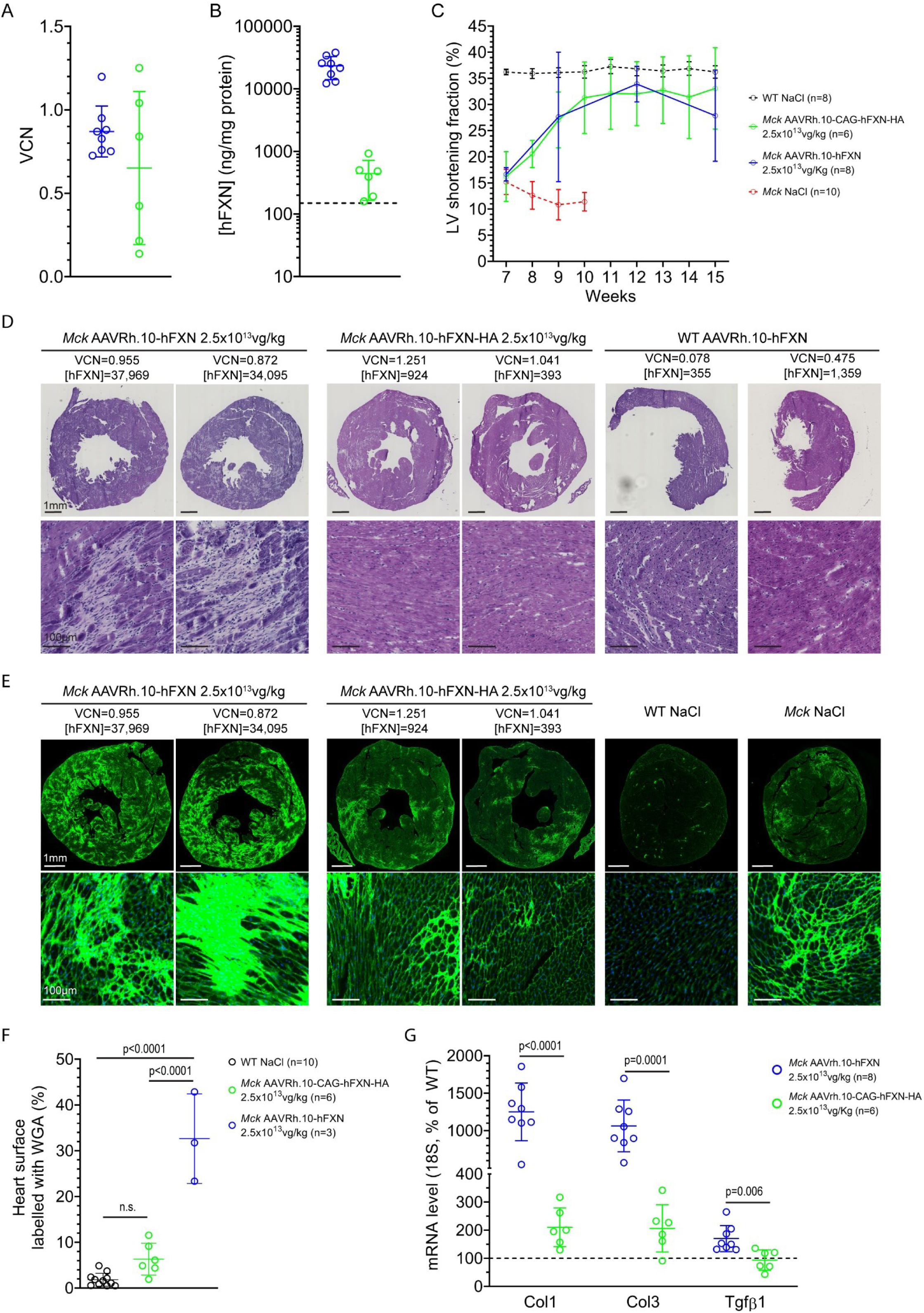
Very high levels of FXN lead to transitory rescue of *Mck* mice cardiac phenotype, followed by acute fibrosis and heart impairment. *Mck* mice were treated at 7-weeks of age with AAVRh.10-hFXN (n=8) or AAVRh.10-CAG-hFXN-HA vector (n=6) at the dose of 2.5×10^13^vg/kg. Data are represented as mean ± SD. **(A)** qPCR quantification of heart VCN. **(B)** ELISA quantification of [hFXN] in the heart, normalized to mg of total heart protein. Blacked-dotted line represents the normal endogenous [mFXN] level in WT mice (147±42 ng/mg). **(C)** Echocardiography assessment of left ventricle (LV) shortening fraction between 7 and 15 weeks of age. For untreated *Mck* mice, historical data were plotted. **(D-E)** Histological analysis of heart fibrosis and cell infiltrates following staining with hematoxylin and eosin (HE) **(D)** or wheat germ agglutinin (WGA, same time exposure). Representative imaging of whole heart tissue section (upper row), with high magnification of LV anterior wall (lower row). Mouse genotype, vector construct and dose, VCN and [hFXN] values are indicated above each panel for all animal. As controls, are reported 9-weeks old untreated *Mck* mice and WT mice treated with NaCl or AAVRh.10-hFXN at 7 weeks and sacrificed at 15 weeks. Scale bars: low magnification 1mm, high magnification 100µm. **(F)** Quantification of heart surface labelled with wheat germ agglutin (WGA) in WT mice (n=10) and the same *Mck* mice treated with AAVRh.10-hFXN (n=3) or AAVRh.10-CAG-hFXN-HA vector (n=6). Statistical significance was evaluated with one-way ANOVA and Turkey post-hoc analysis, p values are reported. **(G)** RTqPCR quantification of Col1a1, Col3a1 and Tgfb1 mRNA level in the heart, normalized to 18S and reported as percentage of WT level. Blacked-dotted line represents normal mRNA level in control mice. Student t-test and p values are reported with n,s p>0.05.

To assess the consequences of FXN overexpression on the therapeutic rescue of *Mck* mice cardiac phenotype, 7-weeks old mice received 2.5×10^13^vg/kg of AAVRh.10-hFXN (n=8) or AAVRh.10-CAG-hFXN-HA vector (n=6). After longitudinal echocardiography evaluation, treated mice were sacrificed at 15 weeks of age. Their respective average heart VCN were very similar (Figure 4A). However, the resulting hFXN levels were much higher in *Mck* mice treated with AAVRh.10-hFXN than with AAVRh.10-CAG-hFXN-HA, with respectively 23,376±9,336 ng/mg and 441±276 ng/mg (Figure 4B). This corresponds to 156-fold and 2.9-fold the normal endogenous FXN level, respectively. Despite this very high overexpression, no significant accumulation of FXN precursor was observed, contrarily to the FXN intermediary form (Figure S12A), suggesting that the majority of the transgenic hFXN protein is targeted to the mitochondria. These data do not support a saturation of the mitochondria import capacity, as a major underlying cause of the mitochondria dysfunction. Moreover, this massive overexpression did not result in the induction of the mitochondrial unfolded protein stress response (mtUPR) (Figure S12B), suggesting that there was no saturation of the chaperon capacity to fold mitochondrial proteins correctly ^35^.

*Mck* mice treated with either vector displayed full correction of the heart function and morphology by 12 weeks of age (Figures 4C and S13A-D). However, the mice treated with the optimized AAVRh.10-hFXN vector started deteriorating afterward, while the mice treated with AAVRh.10-CAG-hFXN-HA displayed sustained full correction (Figures 4C and S13A-D). In line with these results, *Mck* mice treated with the optimized AAVRh.10-hFXN vector displayed substantial increased expression of *Nppa* indicative of pressure/volume overload (Figure S13F), extensive fibrosis and cells infiltrations following histological analysis (Figure 4D-F), which was confirmed by increased expression of genes indicative of on-going fibrosis (Figure 4G), inflammatory response (Figure 5A-B). Like WT mice overexpressing FXN-HA, oxidative stress responsive genes (Hmox1 and Gsta2) were strongly induced (Figure 5C). In contrast, *Mck* mice treated with the non-optimized AAVRh.10-CAG-hFXN-HA vector displayed only few fibrotic patches (Figure 4D-F) and much lower expression for genes involved in fibrosis and inflammatory responses (Figures 4G and 5A).

**Figure 5.**
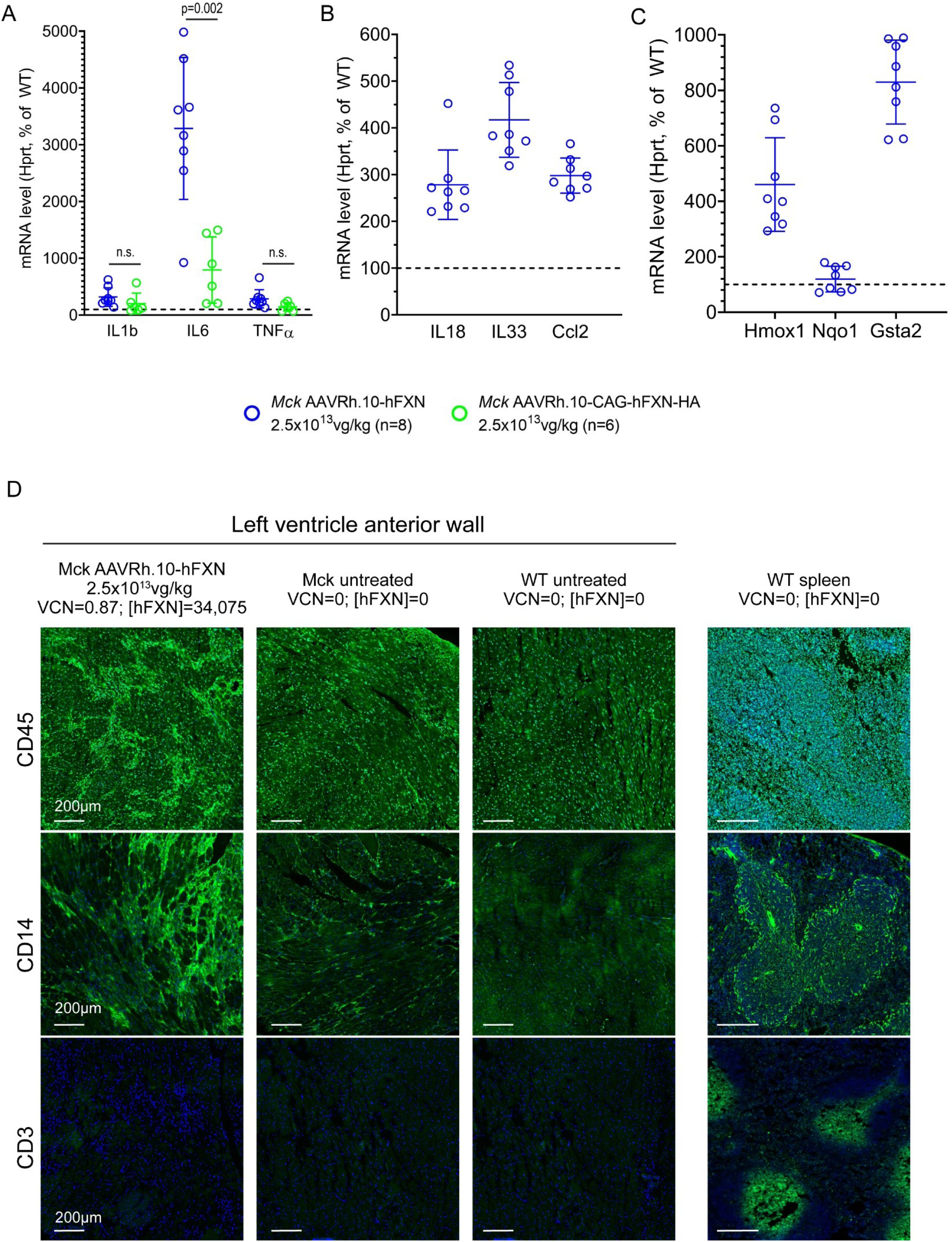
Cardiotoxic overexpression of FXN is associated with acute inflammation but not with adaptative immune response against vector or transgene in *Mck* mice. **(A-C)** *Mck* mice treated at 7-weeks of age with AAVRh.10-hFXN (n=8) or AAVRh.10-CAG-hFXN-HA vector (n=6) at the dose of 2.5×10^13^vg/kg. RTqPCR quantification of gene expression in heart at 15 weeks of age, normalized to Hprt and reported as percentage of WT level. IL1b, IL6 and TNFα **(A)** and IL18, IL33 and Ccl2 **(B)** indicative of inflammatory response. Hmox1, Nqo1 and Gsta2 indicative of oxidative stress response **(C)**. Blacked-dotted line represents normal mRNA levels in control mice. Data are represented as mean ± SD. Student t-test and p values are reported. **(D)** Heart and spleen tissue sections from *Mck* and WT mice treated at 7-weeks of age with vehicle or AAVRh.10-hFXN at the dose of 2.5×10^13^ or 5×10^12^ vg/kg. Immunofluorescent labelling against leucocytes (CD45), monocytes (CD14) and lymphocytes (CD3) cell-type markers. DNA labeled with DAPI. Same time exposure across samples for each labelling. The respective dose, VCN and [hFXN] are reported above each image series. Scale bar, 200µm.

To rule out the potential toxicity intrinsic to the AAVRh.10-hFXN optimized vector, WT mice were also treated at 7 weeks of age and sacrificed at 15 weeks. All of them displayed normal heart histology, with no sign of fibrosis (Figure 4D-E). Furthermore, we examined the liver of the *Mck* mice treated with the optimized AAVRh.10-hFXN vector to determine if there were signs of hepatotoxicity. Despite overexpression up to 97-fold the endogenous FXN level, no obvious liver toxicity was observed (Figure S11E). Altogether, these results rule out a hypothetical intrinsic toxicity of the optimized vector and support the specific relationship between hFXN level and cardiotoxicity. The acute cardiotoxicity was also unlikely to be associated with an immune response against the capsid or the transgene. While we observed significant infiltrations of CD45+ leucocytes and CD14+ monocytes, there was no increase infiltration of CD3+ lymphocytes (Figure 5D). Moreover, the liver did not display any cell infiltrates, which would be expected in the hypothesis of a cytotoxic immune response (Figure S11E). These observations are in line with the observed fibrosis (Figure 4D, E and G) and inflammation (Figure 5A-B), which result from the cardiotoxicity.

The investigation of the mitochondria function and ultrastructure revealed similar alterations as reported above in Mck and WT mice. However, the severity and extend was much higher in mice treated with the optimized vector, in line with the much higher hFXN overexpression across the heart. Histological analysis revealed numerous cardiomyocytes clusters with very high FXN expression and impaired SDH enzymatic activity, covering most of the heart tissue section surface (Figures 6 and S14). In contrast, *Mck* mice treated with the non-optimized vector were rescued for SDH enzymatic activity throughout the heart, except for a few fibrotic patches (Figure S15). Interestingly, the protein level of SDH subunit b (SDHb) was significantly decreased in the heart but not in the liver of *Mck* mice treated with the optimized vector (Figure S10C-E), in line with the apparent lack of liver toxicity despite high levels of overexpression. Furthermore, *Mck* mice treated with the optimized vector also showed impaired NADH enzymatic activity throughout the heart, but normal COX enzymatic activity (Figures 6 and S14), in line with previous observations in WT mice (Figures 1A-S5-S7). In contrast, WT mice treated with the same dose of optimized vector, but overexpressing FXN only by 9.12-fold, displayed normal SDH, NADH and COX enzymatic activity (Figure 6). Altogether, these results confirm the specific correlation between high levels of hFXN protein and the impairment of mitochondrial SDH and NADH enzymatic activities.

**Figure 6.**
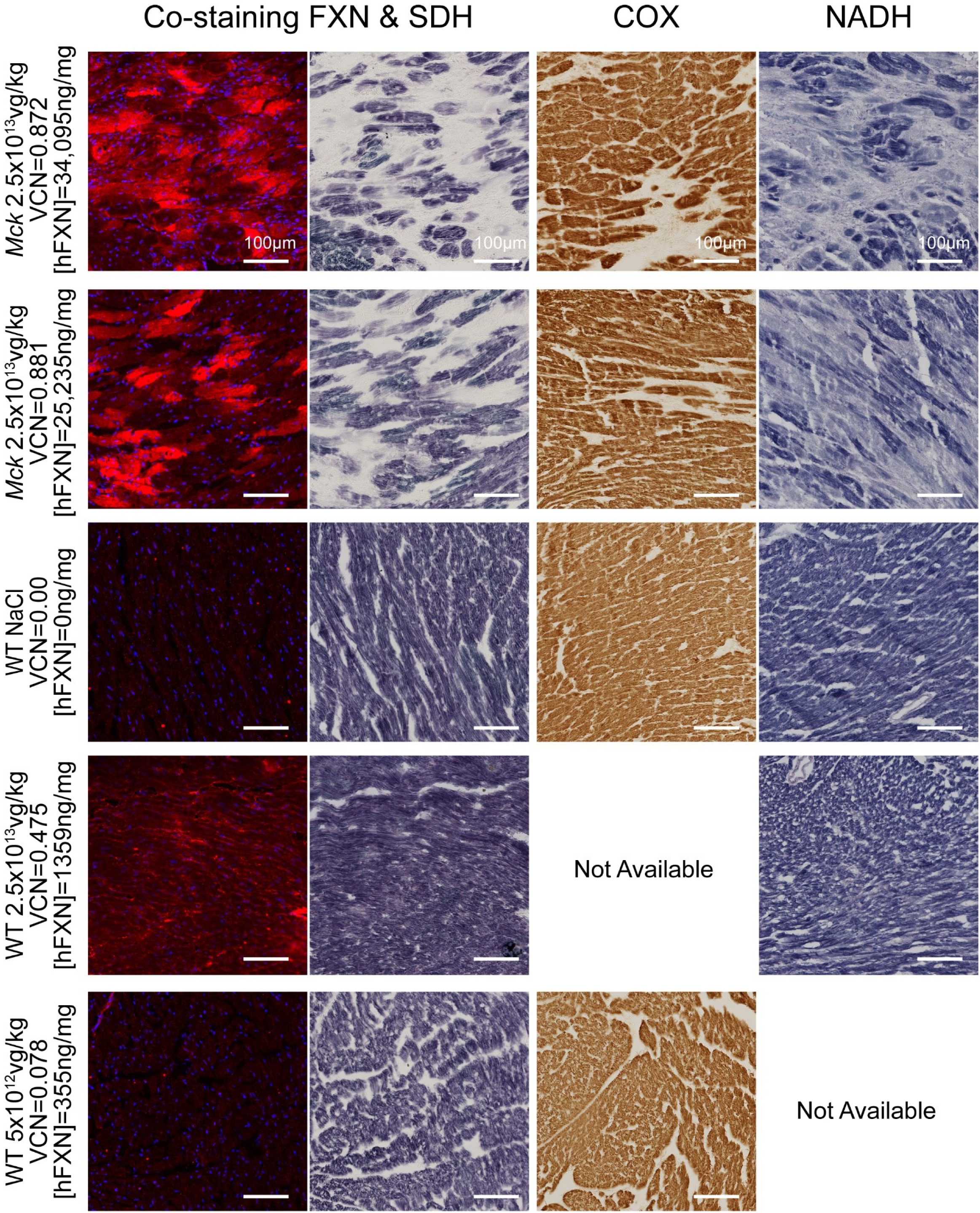
Compromised cardiac gene therapy outcome in *Mck* mice treated with AAVRh.10-hFXN vector is associated with impaired mitochondria respiratory chain function and correlates with the level of FXN overexpression. WT and *Mck* mice treated at 7-weeks of age with vehicle or AAVRh.10-hFXN at the dose of 2.5×10^13^ or 5×10^12^ vg/kg and sacrificed at 15 weeks. Adjacent heart tissue sections were collected to perform histological and colocalization analyses. The two left columns represent the same tissue section and microscopy field co-stained for hFXN by immunofluorescent labelling (constant time exposure) and for succinate dehydrogenase (SDH) activity by in-situ histoenzymatic assay. DNA labeled with DAPI. The middle-right column represents cytochrome C oxidase (COX) hystoenzymatic activity assay. The right-column represents NADH-ubiquinone oxidoreductase (NADH) hystoenzymatic activity assay. DNA labeled with DAPI. The respective dose, VCN and [hFXN] are reported next to each image series. Scale bar, 100μm.

The TEM analysis of the LV myocardium also revealed severe alterations of cardiomyocytes subcellular organization, with scattered and disordered myofibrils, accumulation of swollen mitochondria, with very few visible cristae (Figure 7). Again, we observed electron dense bodies inside the matrix of several mitochondria, which did not appear to be collapsed cristae nor iron deposits. (Figure 7). In line with this observation, *Mck* mice treated with the optimized vector did not display any iron deposits in heart following histological staining (Figure 8).

**Figure 7.**
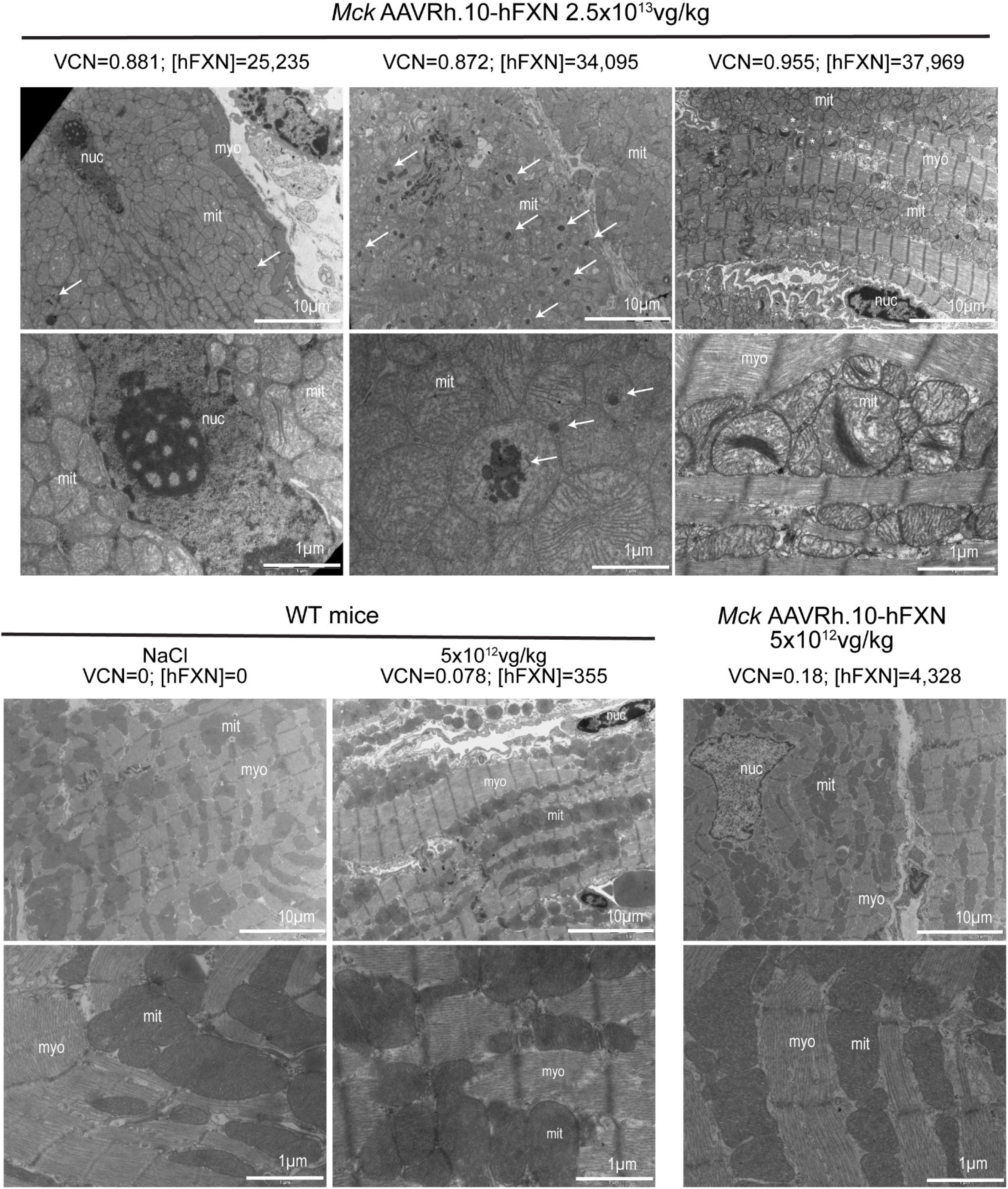
Compromised cardiac gene therapy outcome in *Mck* mice treated with AAVRh.10-hFXN vector is associated with impaired mitochondrial ultrastructure and severe cardiomyocyte disorganization. Transmission electron micrographs from WT and *Mck* mice treated at 7-weeks of age with vehicle or AAVRh.10-hFXN at the dose of 2.5×10^13^ or 5×10^12^ vg/kg and sacrificed at 15 weeks. Representative image of the left ventricle myocardium imaged at low and high magnifications. White arrows indicate mitochondrial electron dense bodies; myo, myofibrils; mito, mitochondria; nuc, nucleus. The respective dose, VCN and [hFXN] are reported for each animal, above each image series. Scale bars: low magnification 10µm, high magnification 1µm.

**Figure 8.**
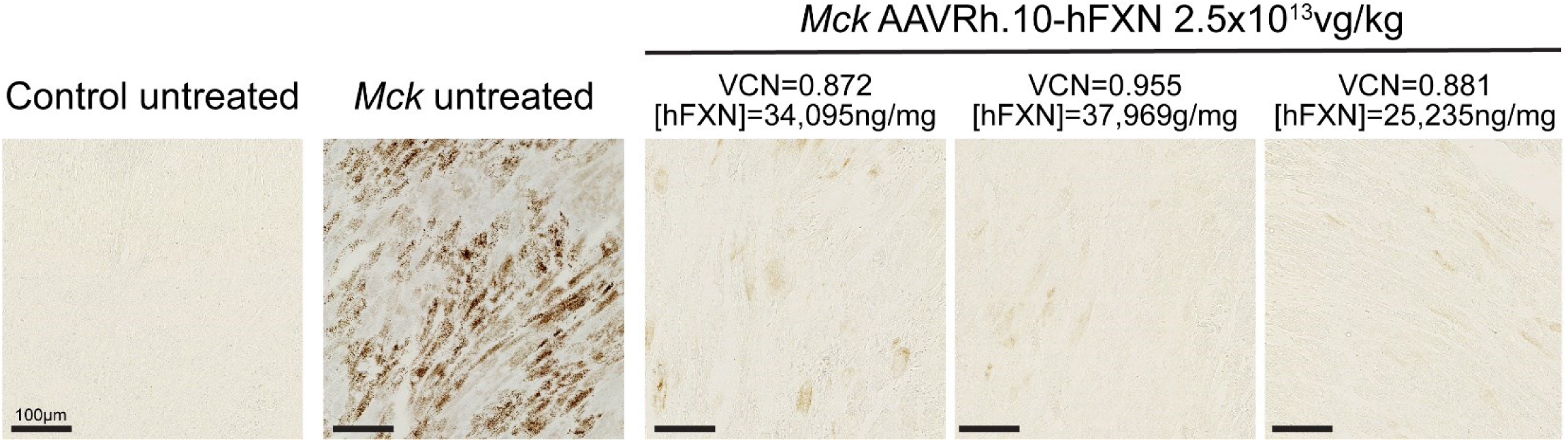
Acute mitochondria and cardiac toxicity caused by very high levels of FXN overexpression is not associated with iron accumulation. WT and *Mck* mice treated at 7-weeks of age with vehicle or AAVRh.10-hFXN at the dose of 2.5×10^13^vg/kg and sacrificed at 15 weeks. Heart tissue sections stained with DAB-enhanced Perls to reveal Fe^3+^ deposits. Representative image of the LV anterior wall. The respective dose, VCN and [hFXN] are reported. Scale bars, 100µm.

## DISCUSSION

Here, we show unequivocally the cardiotoxicity of FXN overexpression when >20-fold the endogenous level and its safety when <9-fold. This was demonstrated in three independent studies, using two different animal models and two different AAVRh.10 vector constructs. The severity of this cardiotoxicity was proportional to the level of FXN overexpression and therefore to the proportion and extent of cardiomyocytes affected throughout the heart. This lead to more or less severe impairment of LV function and morphology, which then evolved either towards compensated heart hypertrophy or severe impairment of heart function. As demonstrated here, this cardiotoxicity might also compromise the therapeutic outcome of cardiac gene therapy for FA. The [hFXN] toxic threshold identified defines more precisely the safe and efficient range for sustained correction of the FA cardiac phenotype.

The cardiac toxic threshold identified, i.e. >9-fold the endogenous level, is in line with previous studies conducted in yeast, mammalian cell lines, and transgenic drosophila and mice models overexpressing constitutively FXN (or Yfh1). Collectively, these studies reported dose depend toxicity when FXN overexpression was higher than 6 to 10-fold the endogenous level ^15-18, 36, 37^, but good tolerance for lower levels ^13, 14, 38-41^. Our results also suggest that this toxicity is not generalizable to all organs and/or cell types, as the mouse liver appeared to be tolerant to FXN overexpression up to 90-fold the normal level, which is acutely toxic to the mouse heart.

Importantly, we ruled out the most obvious alternative hypotheses, including the possible toxicity of a specific vector construct and production lot, vector cytotoxicity or genotoxicity at high dose ^42, 43^, as well as potential immune responses against the vector, transgene and/or transduced cells ^44-47^. Moreover, the mitochondrial capacity to import or process the FXN precursor protein was not impaired, despite very high level of expression.

Cardiotoxic FXN overexpression appeared to be driven by cell autonomous impairment of the mitochondria function and structure, most likely followed by cardiomyocytes cell death, heart fibrosis and LV contractile dysfunction. Strikingly, the mitochondria toxicity affected NADH and SDH enzymatic activities but not COX. This bioenergetics impairment also affected the acetyl-CoA and NAD+/NADH metabolisms. The severe ultrastructure anomalies of the mitochondria most likely contributed to its functional impairment ^21^, which altogether leads to higher mitochondria proliferation and turn-over. These features are reminiscent of the cardiac phenotype of untreated *Mck* mice and are hallmarks of FXN deficient ^7, 9, 12, 29^. However, it is unclear if FXN toxic overexpression was associated or caused by (partial) impairment of Fe-S biosynthesis or handling. Indeed, these two cardiac phenotypes differ from one another by the accumulation of mitochondrial iron upon FXN deficiency ^48^, which was consistently not observed upon FXN overexpressing in the three mouse studies conducted here. These results are in line with previous studies with very high level of FXN overexpression in yeast ^36^, human cells^15^ and drosophila ^17^ where modest increase in labile-iron was shown but no mitochondrial iron deposits. Future studies will be needed to address the underlying mechanism, through time course analysis of pathological events, including exhaustive biochemical analysis of the Fe-S biosynthesis, handling and Fe-S enzymes. The current study also suggests that this toxic threshold was associated with oxidative stress and/or the induction of NRF2-gene responsive genes, as reported in previous studies conducted *in vitro* ^15, 16, 36, 40^and in drosophila ^17^.

These findings would apply to all *in-vivo* AAV-mediated gene replacement strategy for the heart, independently of the AAV serotype or viral vector used. Besides gene replacement strategies, this would also be of concern for alternative approaches if presenting the same risk of acute FXN tissue concentration, either due to their mechanism of action or their delivery modality. This would include synthetic mRNA ^49^, FXN protein replacement ^50, 51^ and *in-vivo* gene transfer of strong artificial transcription factors ^52, 53^. While the current study was focused on the heart, this mitochondrial toxicity is likely to affect other organs. The dorsal root ganglia, the spinal cord, the cerebellum and dentate nucleus, would be of particular interest for future studies, as major tissues of interest for FA neurological therapy. The spleen and kidney are major off-target sites for AAV vectors and should also be considered carefully. Importantly, the FXN level varies largely among these organs ^3, 7^, most likely along cell-type specific metabolism and mitochondria abundance. In present and previous studies ^11, 12^, we have shown that the normal mouse [FXN] level is 149ng/mg in the heart, 49ng/mg in the liver, 12-25ng/mg in DRG and 28ng/mg in the cerebellum. Therefore, we can hypothesize different therapeutic index for each one of these organs. To ensure the safety of future clinical trials, AAV vectors will need to be designed to manage appropriately their expression profile across these different tissues. This could be achieved with (1) synthetic weak promoter or the endogenous human frataxin promoter ^54, 55^, (2) vector de-targeting strategies^56, 57^, and/or (3) the design of expression cassette responsive to negative cellular feedback loop. Leveraging our knowledge of the cellular response to toxic FXN overexpression might help achieve the latter, for example with 3’UTR target sequence in the vector specific to endogenous microRNA induced by the ISR ^58, 59^ or NRF2-mediated oxidative stress response ^60, 61^. In addition, specific routes of drug delivery might also present risk of high local concentration, such as intracardiac injection ^62^ or local delivery in the central nervous system ^63, 64^. Due diligence in the design of appropriate GLP pharmacokinetic and toxicity studies, in relevant large animal model, will be crucial for the development of safe clinical therapeutic protocol. Finally, the identification of biomarkers to monitor subclinical toxicity and manage such event, would be very advantageous. In this regard, the current study has identified potential candidates, including FGF21 and LGASL3, that will be explored in future studies.

## CONCLUSION

In summary, the overexpression of FXN is safe when ≤ 9-fold the normal endogenous level but cardiotoxic when greater than 20-fold. The pathological mechanism was partially elucidated and is most likely primed by the alteration of the mitochondria structure and function. Depending on the percentage of cardiomyocytes affected, this resulted either in subclinical or acute cardiotoxicity. The toxic threshold and potential read-out for toxicity identified here will support the design of robust and meaningful GLP toxicity studies, for safer FA gene therapy clinical trials.

## ACKNOWLEDGMENTS

We thank Dr Hugues Jacob, Karim Hnia, Nicolas Charlet-Berguerand and Pr. Jean-Louis Mandel for scientific discussions and Dr Wyatt Delphino and Mehdi Gasmi for critical review of the manuscript. We thank Pr. Ronald G. Crystal (Department of Genetic Medicine, Weill Cornell Medicine, New York) for the production of the optimized construct and vector, Dr. Mustapha Oulad-Abdelghani for providing the 4F9 mouse monoclonal antibodies against FXN. We thank Dr. Tania Sorg, Ghina Bou-About, Mrs Josiane Hergueux, the mouse clinic institute phenotyping platform and Pr. Luc Dupuis for logistical and technical assistance. This work was supported by the Lefoulond Delalande foundation - Institut de France [post-doctoral research fellowships years 2016]; the Friedreich Ataxia Research Alliance [Keith Michael Andrus Cardiac Research Award year 2016]; the French Muscular Dystrophy Association AFM-Téléthon [Research grant year 2014); AAVLIFE SAS [sponsored research agreement years 2014-2016]. This study was also supported by the grant ANR-10-LABX-0030-INRT, a French State fund managed by the Agence Nationale de la Recherche under the frame program Investissement d’Avenir ANR-10-IDEX-0002-02.

## AUTHORS CONTRIBUTION

B. Belbellaa and H. Puccio conceived the project. B. Belbellaa conducted and analyzed the experiments. L. Reutenauer managed mice production. N. Messaddeq performed transmitted electron microscopy. L. Monassier provided logical support for echocardiography. B. Belbellaa and H. Puccio wrote the manuscript.

## SUPPLEMENTAL DATA ITEMS

**Figure S1.**
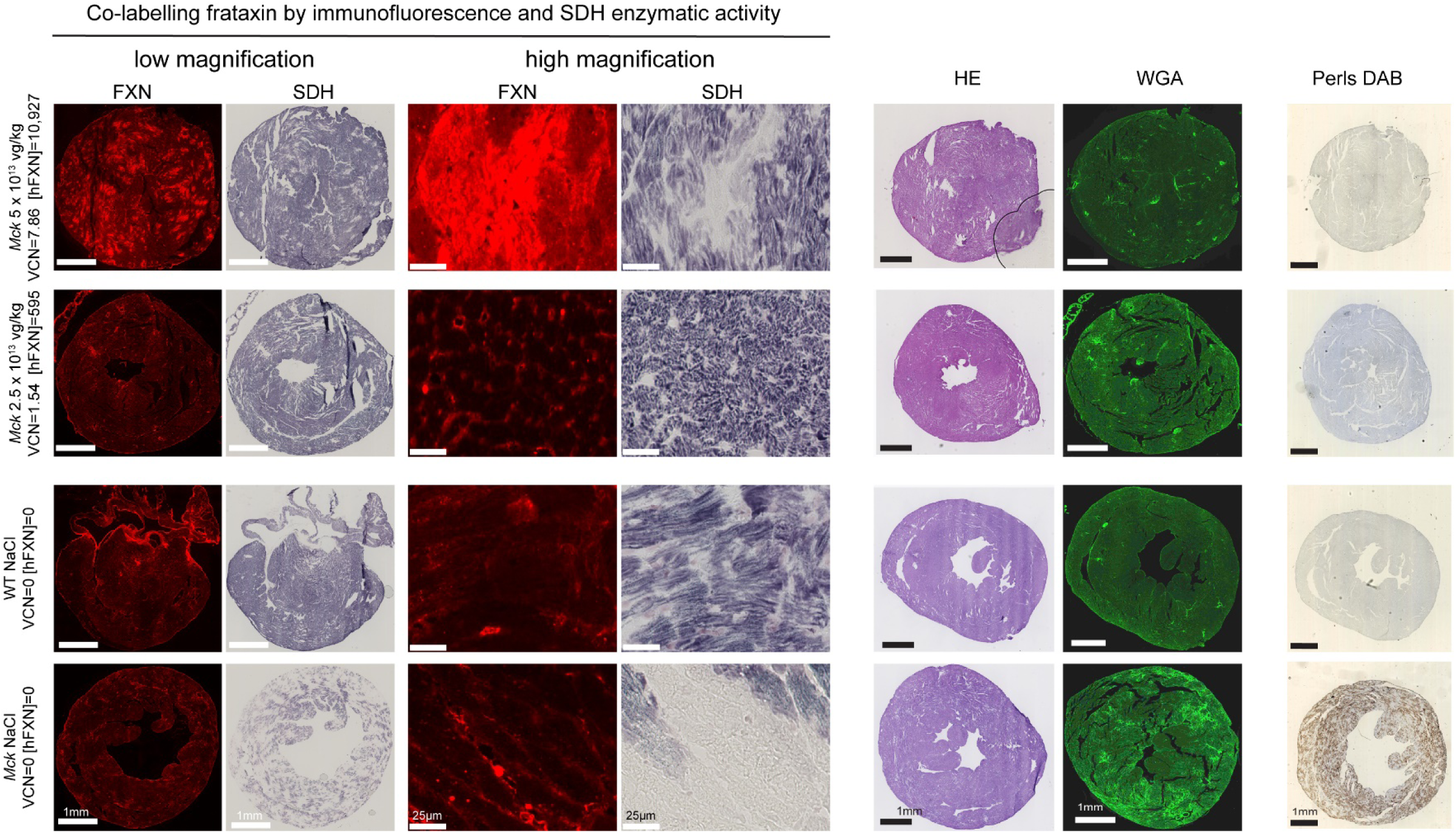
*Mck* mice cardiomyocytes overexpressing FXN-HA at 73-fold the endogenous level display impaired succinate dehydrogenase enzymatic activity without heart fibrosis nor iron cell deposits. Adjacent heart tissue sections from WT and *Mck* mice treated at 5 weeks of age with NaCl or AAVRh.10-CAG-hFXN-HA vector at doses of either 5×10^13^ or 2.5×10^13^ vg/kg and sacrificed at 12 weeks. 9 weeks old untreated *Mck* mice were used as control. The four columns on the left represent the same tissue sections and microscopy fields co-stained for hFXN by immunofluorescent labelling (same time exposure) and for succinate dehydrogenase (SDH) activity by *in-situ* histoenzymatic assay. Scale bars: low magnification 1mm, high magnification 25µm. The two middle-right columns correspond to histological analysis of heart fibrosis and cell infiltrates following H&E or WGA staining (same time exposure). The last column on the right represents DAB-enhanced Perls labelling of iron deposits with methyl green counterstaining. Scale bars, 1mm. The corresponding vector copies per diploid genome (VCN) and tissue concentration in human FXN ([hFXN]) are reported next to each image series.

**Figure S2.**
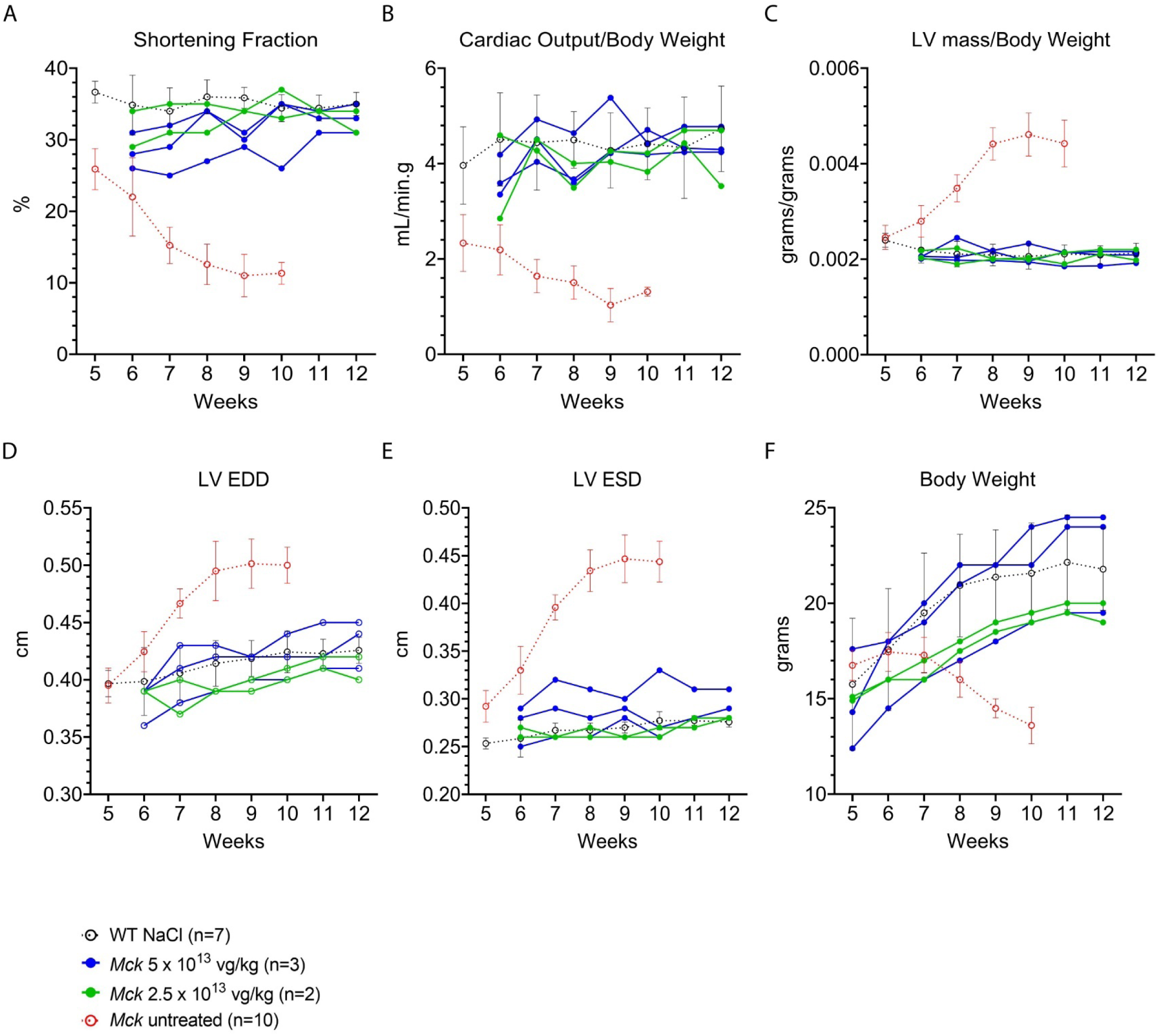
Longitudinal echocardiography evaluation of *Mck* mice overexpressing FXN-HA up to 73-fold the normal level did not reveal subpar rescue of heart function or morphology. *Mck* mice treated at 5 weeks of age with AAVRh.10-CAG-hFXN-HA vector at the dose of 5×10^13^ (n=3) or 2.5×10^13^ (n=2) vg/kg are represented as individual kinetics. Data are represented as mean ± SD for WT control mice (n=7) and untreated *Mck* mice (n=10). For untreated *Mck* mice, historical data were plotted. Statistical analyses are reported in Table S1. **(A)** Left ventricle (LV) shortening fraction. **(B)** Cardiac blood output measured at the aorta (CO) and normalized to body weight (BW). **(C)** LV mass normalized to BW. **(D)** LV end-diastole diameter (EDD). **(E)** LV end-systole diameter (ESD). **(F)** Body weight. Figure adapted from belbellaa et al ^1^.

**Figure S3.**
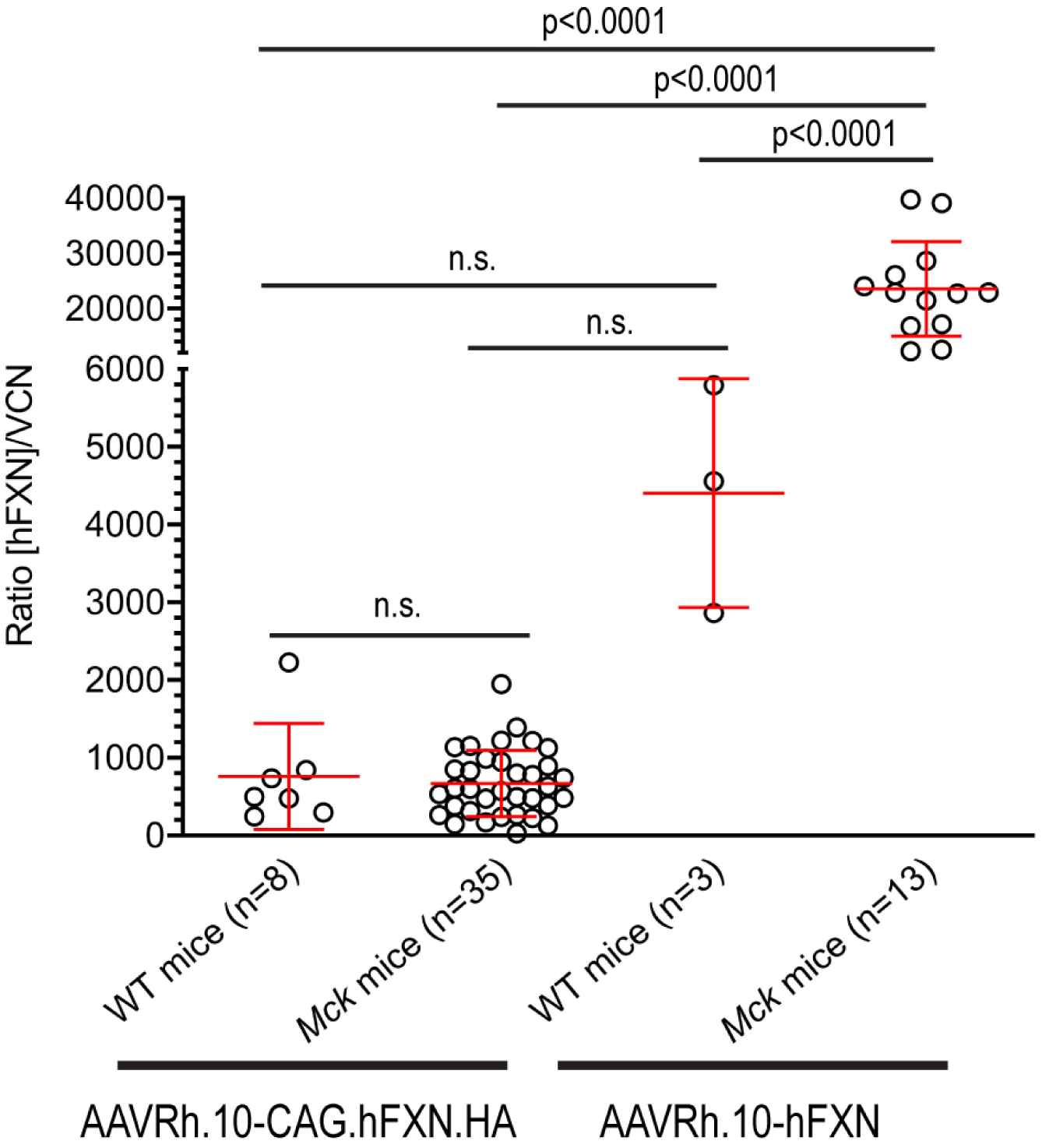
Comparison of AAVRh.10 vectors potency in the heart of WT and *Mck* mice treated at 7 weeks and sacrificed at 15 weeks. Vector potency was quantified in each individual mouse heart where the tissue concentration of FXN protein was measured by ELISA assay (expressed as ng of FXN per mg of total protein) and normalized by the average vector DNA copies per diploid genome. Data are reported as mean ± SD. One-way ANOVA statistical analysis, p values are reported with n.s. p>0.05.

**Figure S4.**
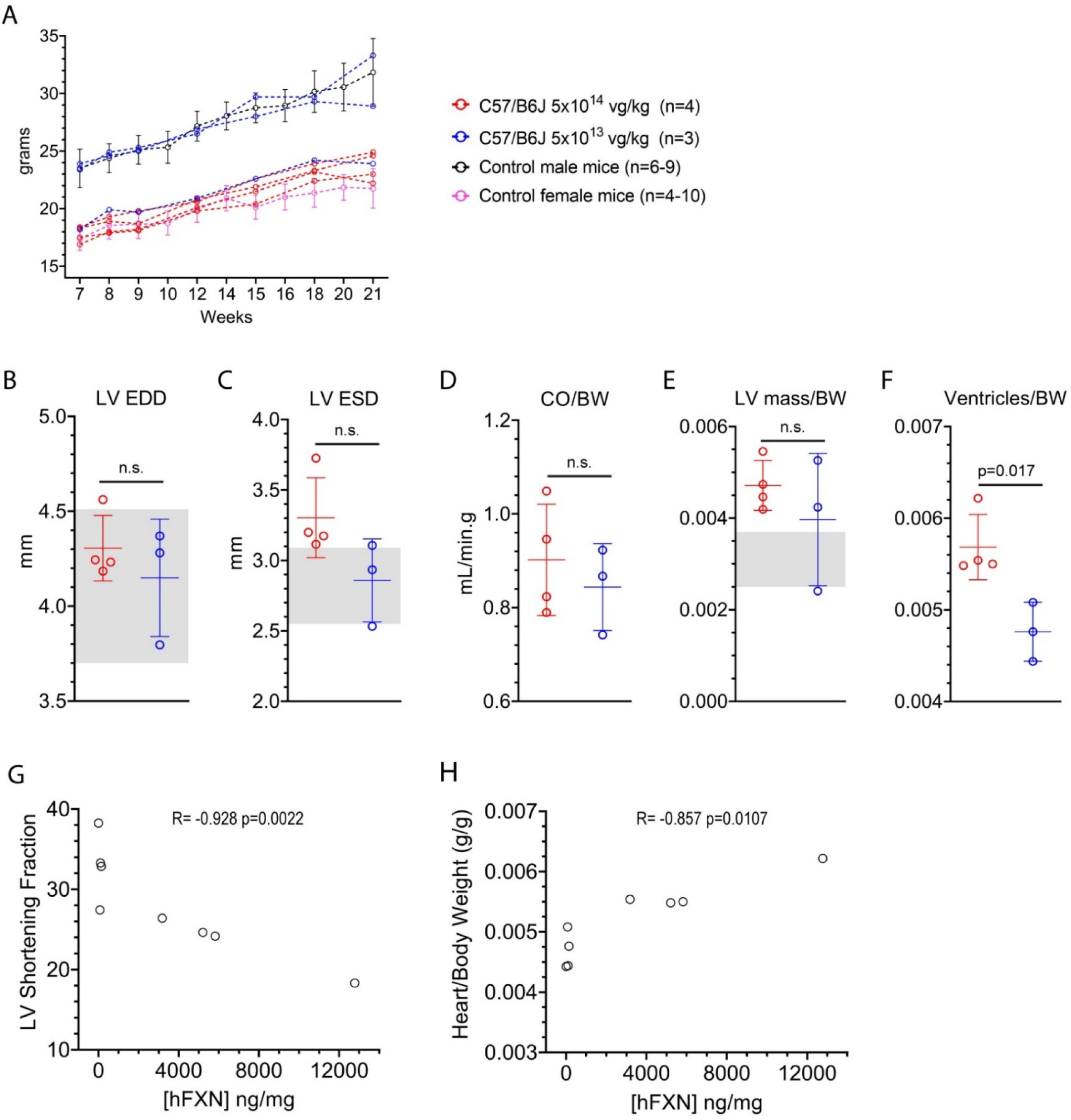
Detailed echocardiography analysis of WT C57/B6J mice treated with AAVRh.10-CAG-hFXN-HA vector and overexpressing FXN-HA. (**A**) Body weight reported as individual mouse kinetic with males in blue and females in red. For control mice, historical data were plotted as mean ±SD. (**B-E**) Echocardiography analysis of left ventricle (LV) function and morphology at 21 weeks of age. Data are reported as individual datapoints with mean ±SD. Grey-shaded area corresponds to normal values range. Student t test and p values are reported. **(B)** LV end-diastole diameter (EDD). **(C)** LV end-systole diameter (ESD). (**D**) Cardiac blood output measured at the aorta (CO) normalized to body weight (BW). **(E)** LV mass normalized to BW. (**F**) Heart ventricles weight measured upon necropsy, normalized to BW. (**G-H**) Regression analysis between heart tissue concentration [hFXN] and LV shortening fraction (**G**) or with heart ventricle weight normalized to BW (**H**). Spearman non-parametric correlation coefficient and p value are reported.

**Figure S5.**
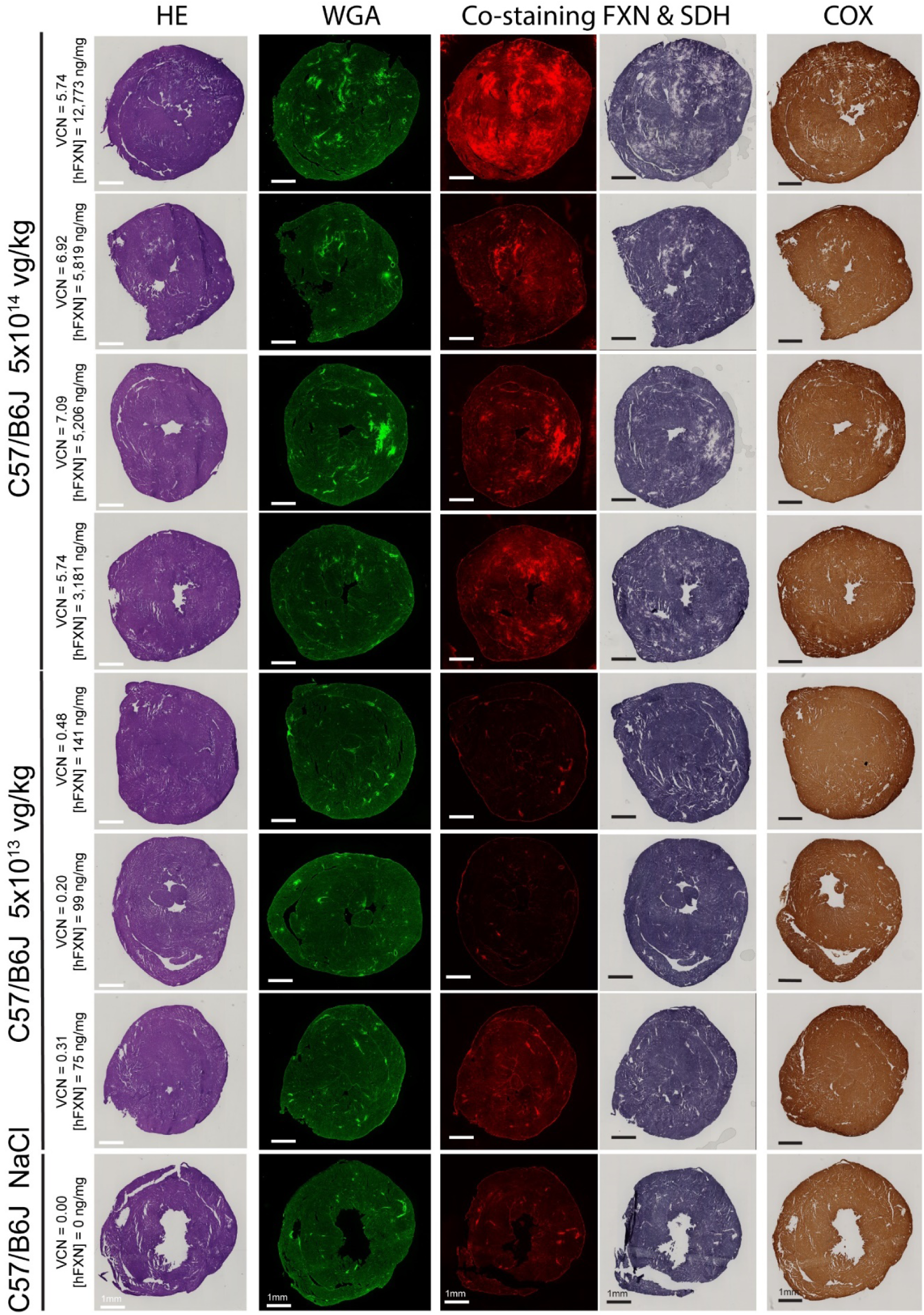
Detailed heart histology analysis of WT C57/B6J mice overexpressing FXN-HA protein. Wild-type (WT) C57/B6J mice were treated at 7-weeks of age with vehicle or AAVRh.10-CAG-hFXN-HA vector at the dose of 5×10^13^ (n=3) or 5×10^14^ (n=4) vg/kg, and then sacrificed at 21 weeks of age. Adjacent heart tissue sections were collected to perform histological analysis. The two left columns correspond to histological analysis of heart fibrosis and cell infiltrates following H&E or WGA staining. The two right-middle columns represent the same tissue section and microscopy field co-stained for FXN by immunofluorescent labelling and for succinate dehydrogenase (SDH) activity by in-situ histoenzymatic assay. The right column represents cytochrome c oxidase (COX) histoenzymatic activity assay. The corresponding dose, VCN and [hFXN] are reported next to each image series. Scale bar, 1mm. Same time exposure used for each labelling series.

**Figure S6.**
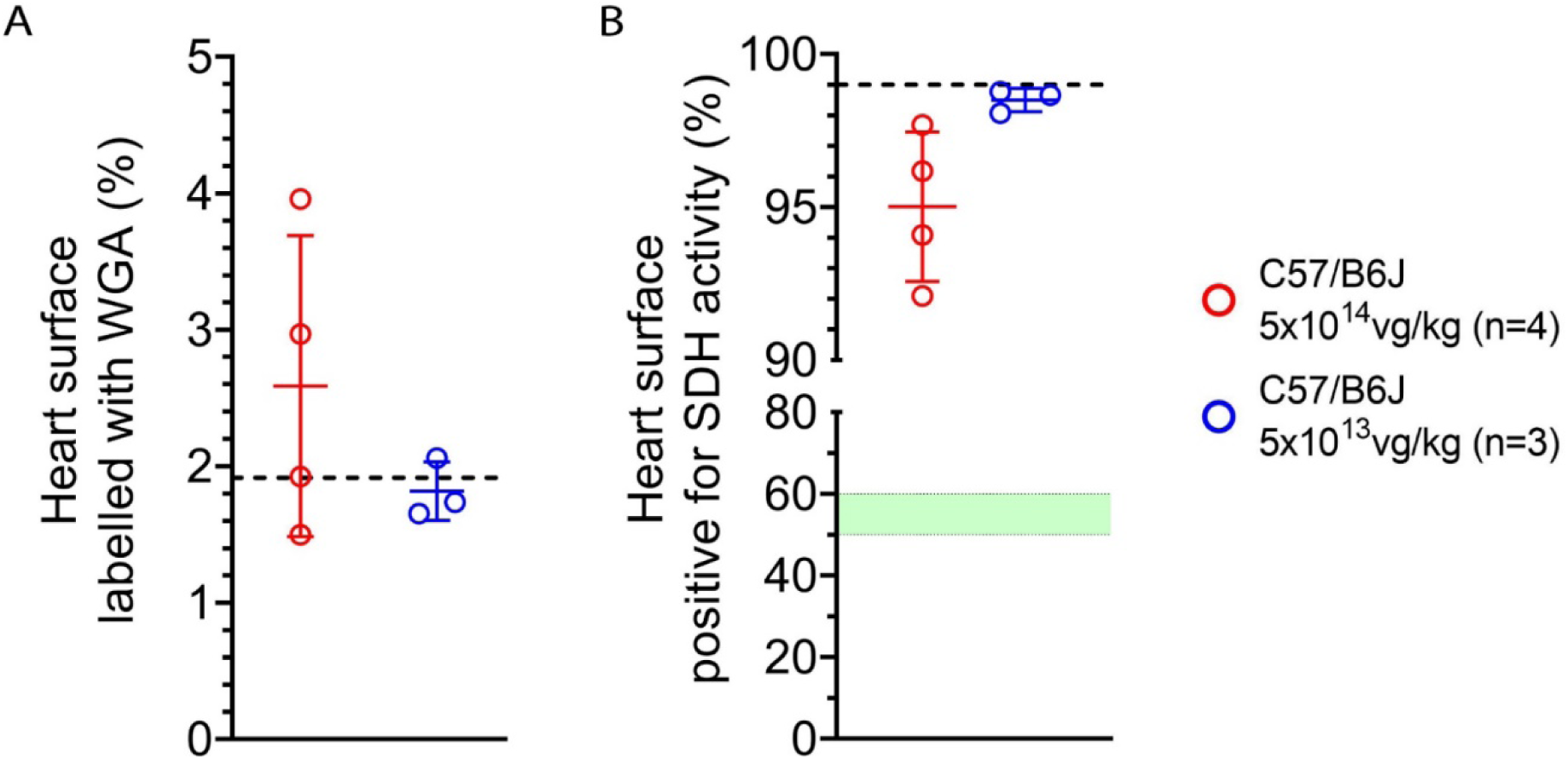
Subclinical cardiotoxic overexpression of FXN-HA in WT C57/B6J mice is associated with overall sparse fibrosis and mitochondria function impairment across the heart. **(A)** Quantification of heart surface labelled with wheat germ agglutin (WGA). **(B)** Quantification of heart surface positive for succinate dehydrogenase (SDH) enzymatic activity.

**Figure S7.**
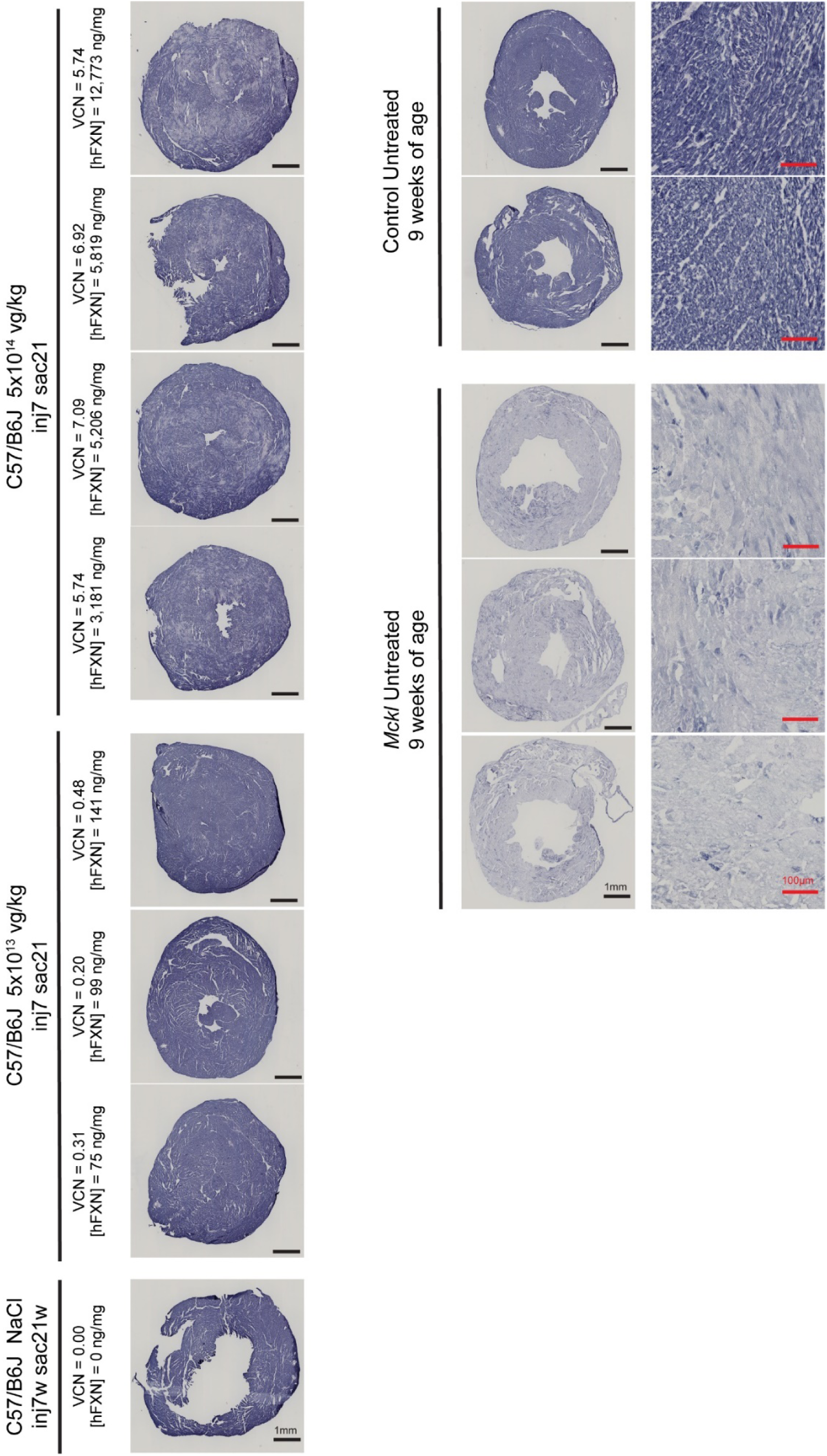
NADH-ubiquinone oxidoreductase (NADH) enzymatic activity is severely reduced throughout the heart of untreated *Mck* mice heart and to a much lesser extend in WT C57/B6J mice overexpressing ≥ 5,206ng of FXN per mg total heart protein. *In-situ* enzymatic activity assay on heart tissue sections from untreated *Mck* and WT C57/B6J and treated WT C57/B6J. The corresponding AAVRh.10-CAG-hFXN-HA vector dose, VCN and [hFXN] are reported next to each image. Scale bar: low magnification, 1mm; high magnification, 100µm.

**Figure S8.**
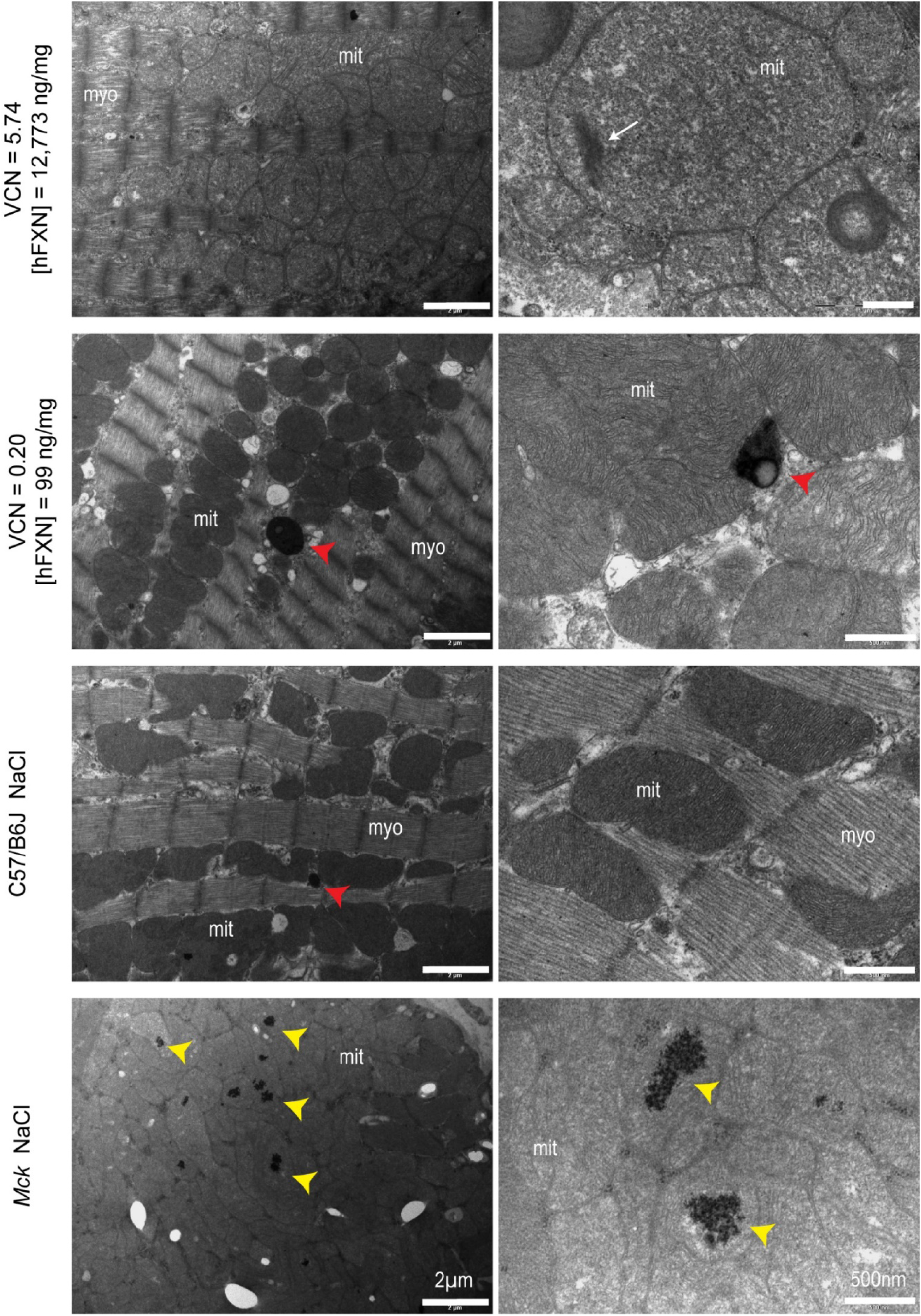
Transmitted electron microscopy imaging of heart myocardium following staining of ferritin. Ferritin labelled with bismuth sodium tartrate in heart ultra-thin sections from WT mice treated with vehicle or AAVRh.10-CAG-hFXN-HA vector at the dose of 5×10^14^ (n=4) or 5×10^13^ (n=3) at 7 weeks of age and sacrificed at 21 weeks. Heart tissue sections from untreated *Mck* mice sacrificed at 7 weeks of age were used as controls. Red arrowheads indicate lysosome containing ferritin; yellow arrowheads indicate ferritin inside mitochondria; white arrows indicate mitochondrial electron dense bodies; myo, myofibrils; mito, mitochondria. Left column represents low magnifications (Scale bar, 2µm) and right column high magnification (500nm). The respective dose, VCN and [hFXN] are reported next to each image series.

**Figure S9.**
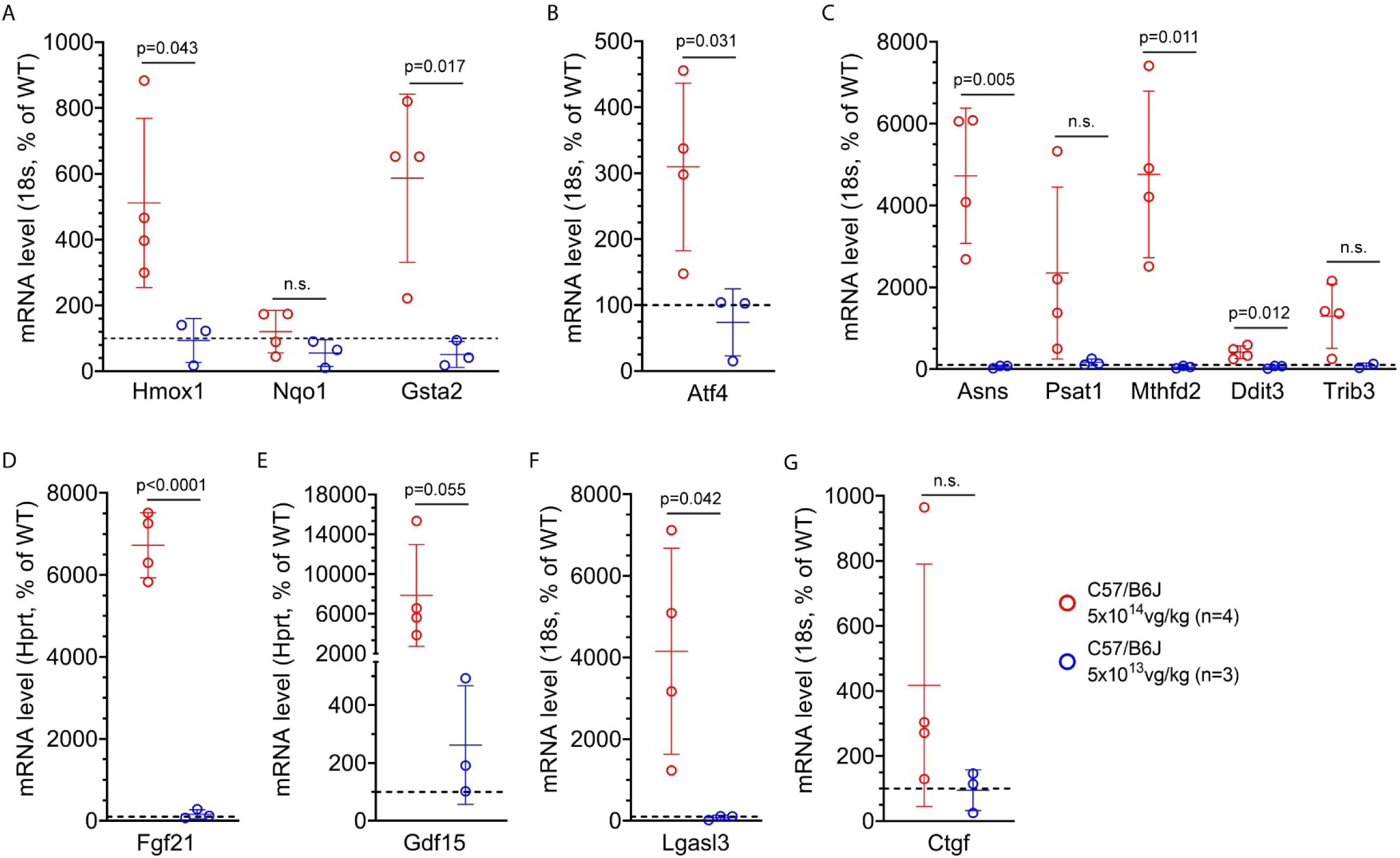
Cardiotoxic FXN overexpression is associated with the induction of the oxidative stress response and the integrated stress response. RTqPCR analysis of heart gene expression in WT C57/B6J mice treated with 5×10^13^ or 5×10^14^ vg/kg. mRNA levels are reported as mean ± SD, normalized to 18S or Hprt and as percentage of WT level. Blacked-dotted line corresponds to control level. Student t-test and p values are reported. **(A)** NRF2-responsive genes indicative of oxidative stress response. (**B-C**) Integrated stress response (ISR) induced genes. **(D-G)** ISR-target genes encoding for proteins secreted in plasma and previously identified as biomarkers of cardiac or mitochondrial dysfunction.

**Figure S10.**
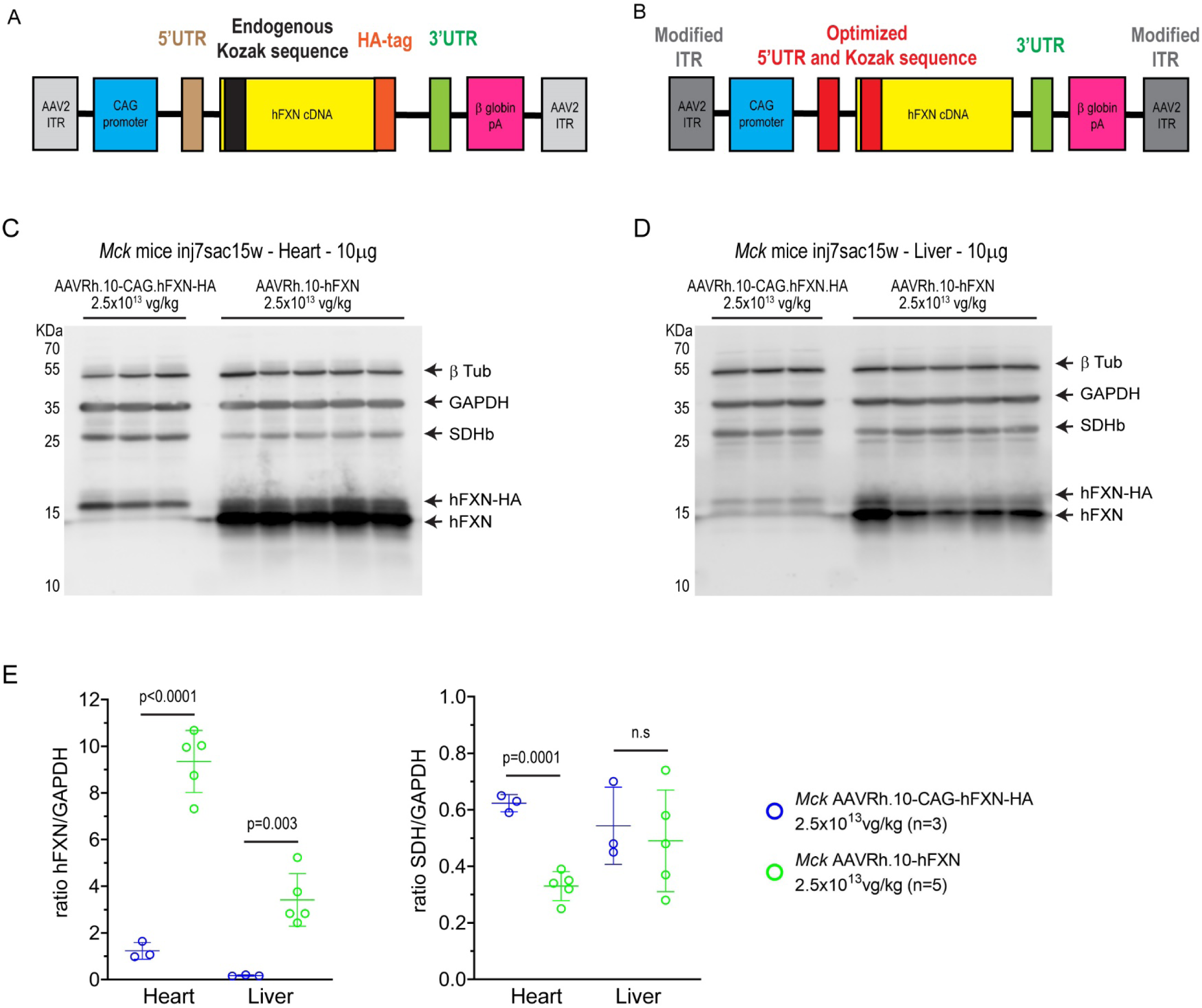
Description of AAVRh.10 vector constructs encoding human FXN and comparison of vector expression and potency in heart and liver of WT and *Mck* mice. **(A)** AAVRh10-CAG-hFXN-HA vector construct, used in the proof of concept and dose response mouse studies ^1, 2^. **(B)** AAVRh.10-hFXN optimized vector construct. **(C-D)** Western blot analysis comparing the expression of AAVRh.10 vectors in the heart **(C)** and liver **(D)** of *Mck* treated with 2.5×10^13^ vg/kg at 7 weeks and sacrificed at 15 weeks. Electrophoresis of 10µg total protein on SDS PAGE 14% gel. Immunoblotting against both human (hFXN) and mouse (mFXN) frataxin, Complex II 30KDa subunit (SDHb), Glyceraldehyde-3-phosphate dehydrogenase (GAPDH) and beta-tubulin (β-Tub). hFXN and hFXN-HA migrate at sensibly different molecular weights. **(E)** Quantification of the relative protein levels of hFXN and SDH, normalized to GAPDH, in the heart and liver. Student t-test, p values are reported, not significant ns>0.05.

**Figure S11.**
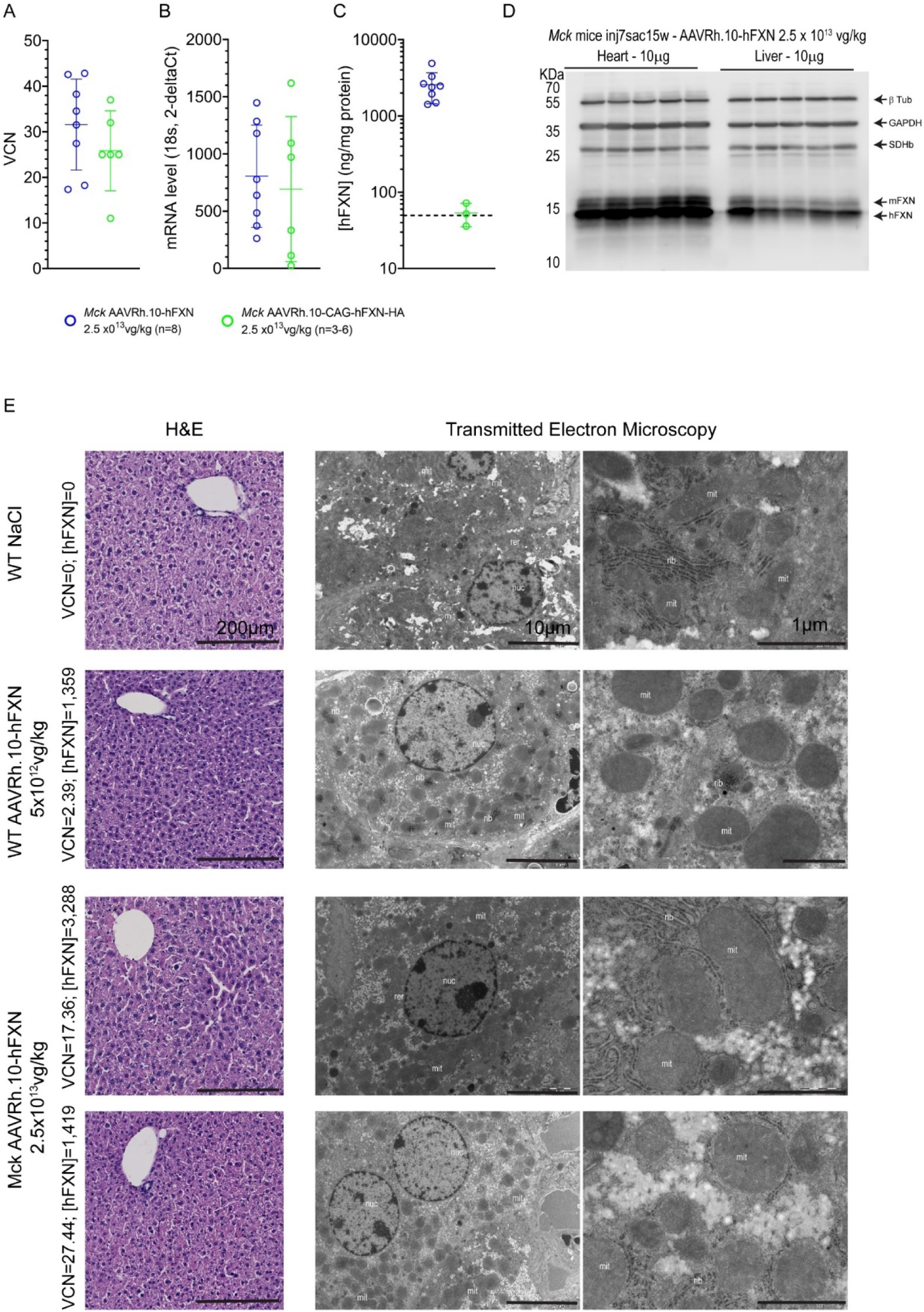
Liver overexpression up to 87-fold the normal level does not result in impaired FXN protein processing, acute liver toxicity or alteration of hepatocytes and mitochondria ultrastructure. **(A)** qPCR analysis of vector copies per diploid genome in liver. Student t-test: p=0.2833. **(B)** RTqPCT analysis of hFXN mRNA level normalized to 18s and reported as 2^-deltaCt^. Student t-test: p=0.06994. **(C)** ELISA quantification of [hFXN] reported as ng per mg of total liver protein. Black dotted line represents, normal endogenous level of mFXN in WT C57/B6J mice liver (n=5), i.e. 49.6±7.8 ng per mg of protein. Student t-test: p=0.0046. Data are reported as mean ± SD. **(D)** Comparative analysis of AAVRh.10-hFXN vector expression in the heart and liver of the same *Mck* mice (n=5) treated with 2.5×10^13^ vg/kg at 7 weeks and sacrificed at 15 weeks. Electrophoresis of 10µg total protein on SDS PAGE 14% gel. Immunoblotting against both human (hFXN) and mouse (mFXN) frataxin, Complex II 30KDa subunit (SDHb), Glyceraldehyde-3-phosphate dehydrogenase (GAPDH) and beta-tubulin (β-Tub). **(E)** Liver tissue sections from WT and *Mck* mice treated with NaCl or AAVRh.10-hFXN at 7 weeks and sacrificed at 15 weeks. The corresponding dose, VCN and [hFXN] are reported next to each image series. Left column represents hematoxylin and eosin staining of 10µm thick section. The two-right columns represent transmitted electron microscopy observation of ultra-thin sections (Scale bars: left, 10µm; right, 1µm).

**Figure S12.**
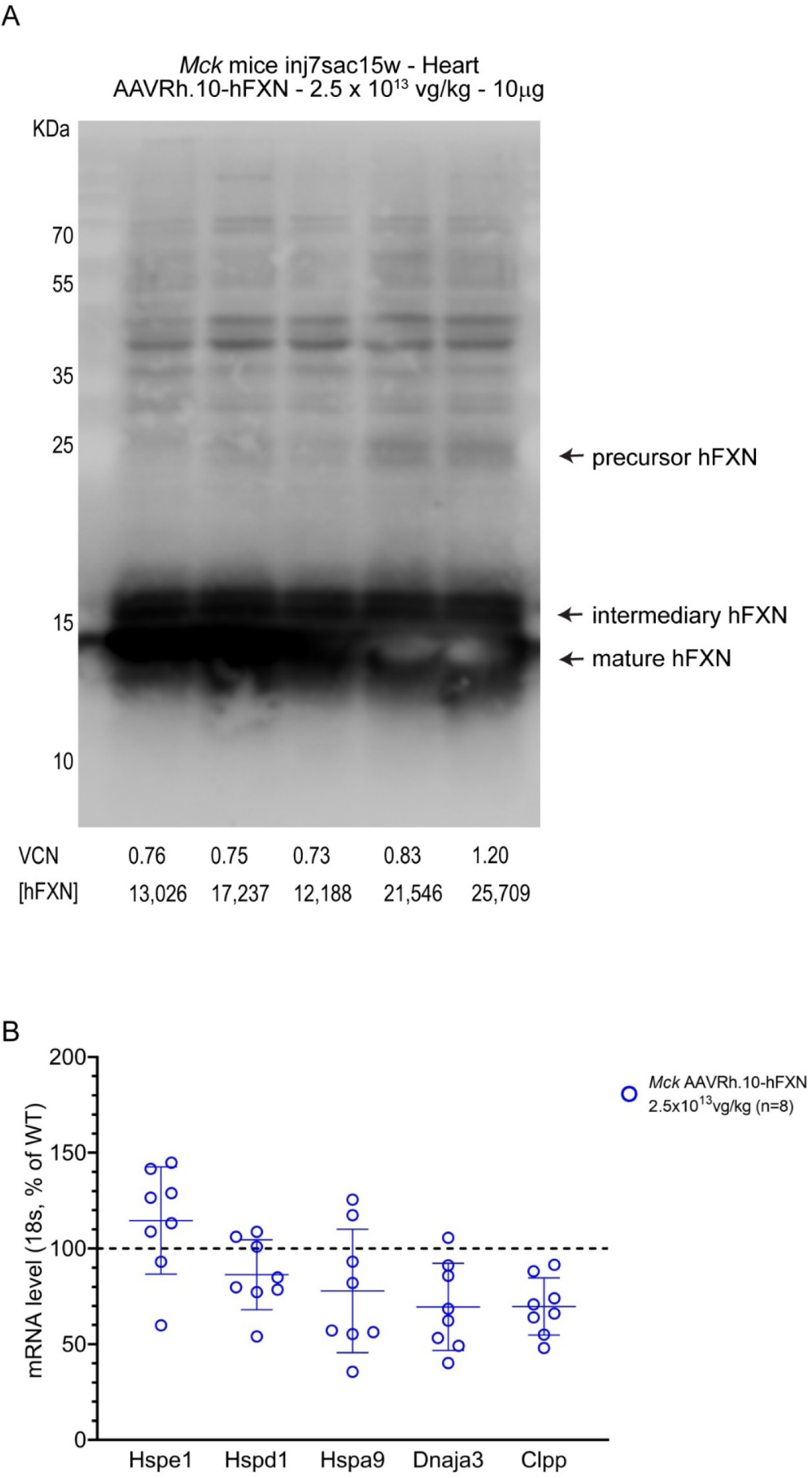
Cardiac overexpression up to 172-fold the normal level does not lead to FXN precursor accumulation but to some extend of the intermediary form, without induction of the mitochondrial unfolded protein stress response (mtUPR). **(A)** WB analysis of total heart protein heart extract (10µg per well) from *Mck* mice treated with 2.5×10^13^vg/kg of AAVRh.10-hFXN at 7 weeks of age and sacrificed at 15 weeks. The respective VCN and [hFXN] are reported. Immunoblotting against FXN, with detection of the mature hFXN at 14KDa, the intermediate form at 18KDa and the precursor form at 25KDa. 10µg of protein were loaded per well in SDS PAGE 14% gel. **(B)** RTqPCR analysis of mtUPR-responsive genes, mRNA levels are normalized to 18s and reported as percentage of control level. Black-dotted line represents control level.

**Figure S13.**
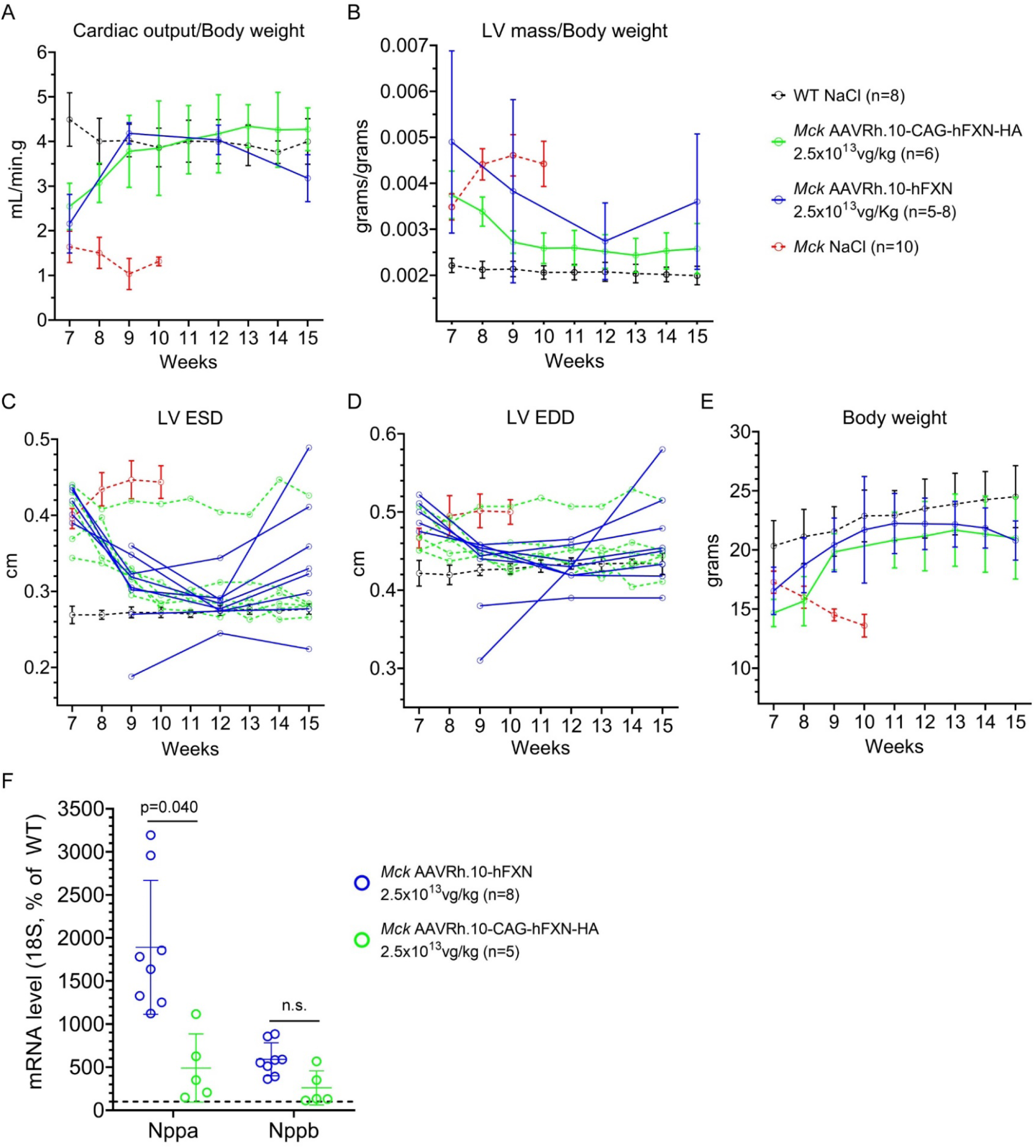
Detailed echocardiography analysis of *Mck* mice treated at 7 weeks of age with 2.5×10^13^ vg/kg of AAVRh.10-CAG-hFXN-HA or AAVRh.10-hFXN vector. For untreated WT and *Mck* mice, historical data were plotted. Data are represented as mean ± SD. Statistical analysis is reported in Table S2. **(A)** Cardiac blood output measured at the aorta, normalized to body weight. **(B)** LV mass normalized to body weight. **(C)** LV end-systole diameter (LV ESD). **(D)** LV end-diastole diameter (LV EDD). **(E)** Body weight. **(F)** RTqPCR analysis of Nppa and Nppb gene expression in the heart. Student bilateral t-test and p values are reported.

**Figure S14.**
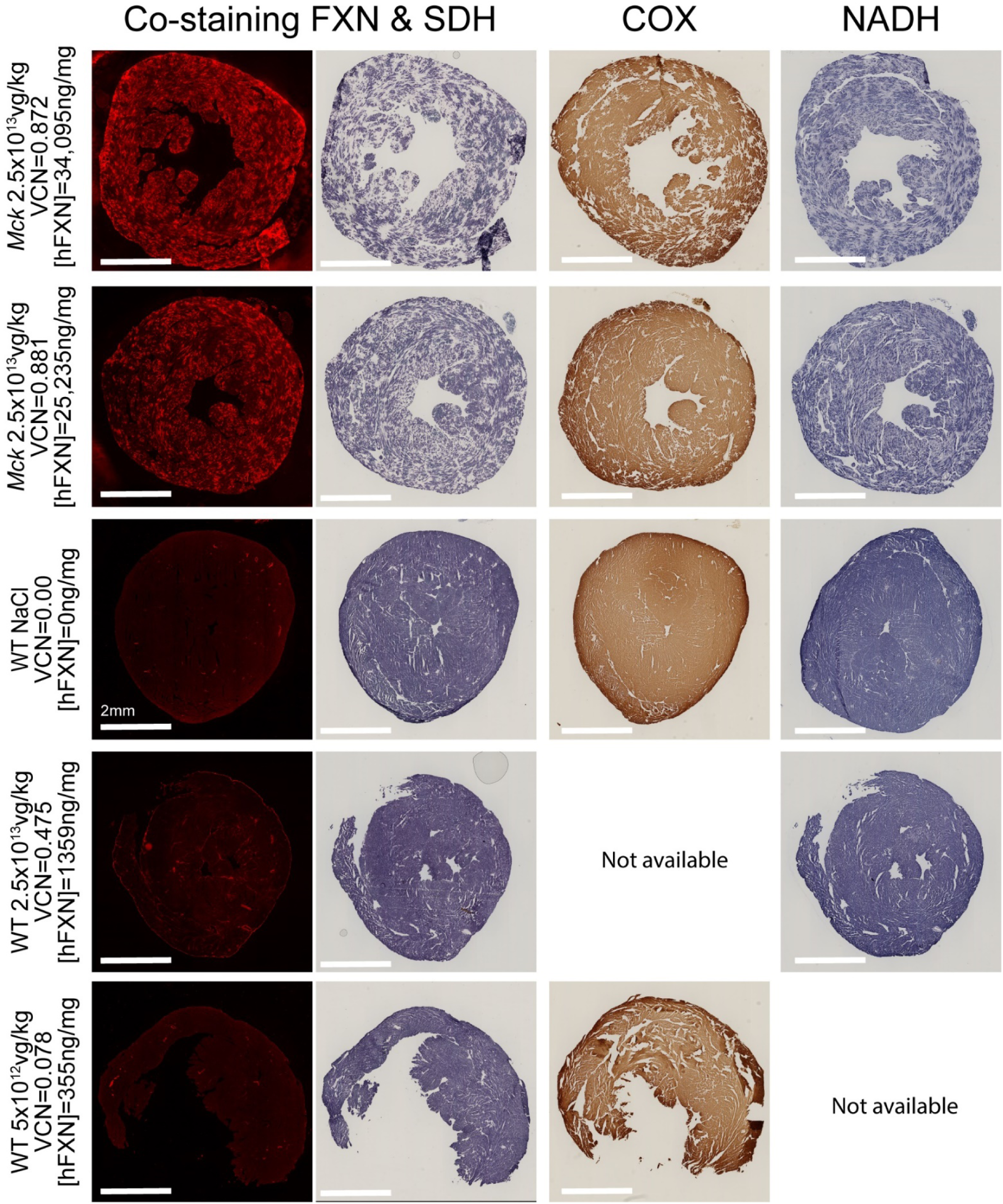
Extended heart histology analysis of FXN overexpression and mitochondria respiratory chain function in *Mck* mice treated at 7 weeks with 2.5×10^13^vg/kg and sacrificed at 15 weeks. Adjacent heart tissue sections were collected to perform co-labelling of hFXN and SDH *in-situ* histoenzymatic assay (two-left columns), *in-situ* histoenzymatic assay for COX (middle-right column) or for NADH (right column). As controls, WT mice were treated with vehicle or AAVRh.10-hFXN vector at the doses of 2.5×10^13^ or 5×10^12^ vg/kg. The respective dose, VCN and [hFXN] are reported above each image series. Scale bar, 2mm. Same time exposure for each labelling series.

**Figure S15.**
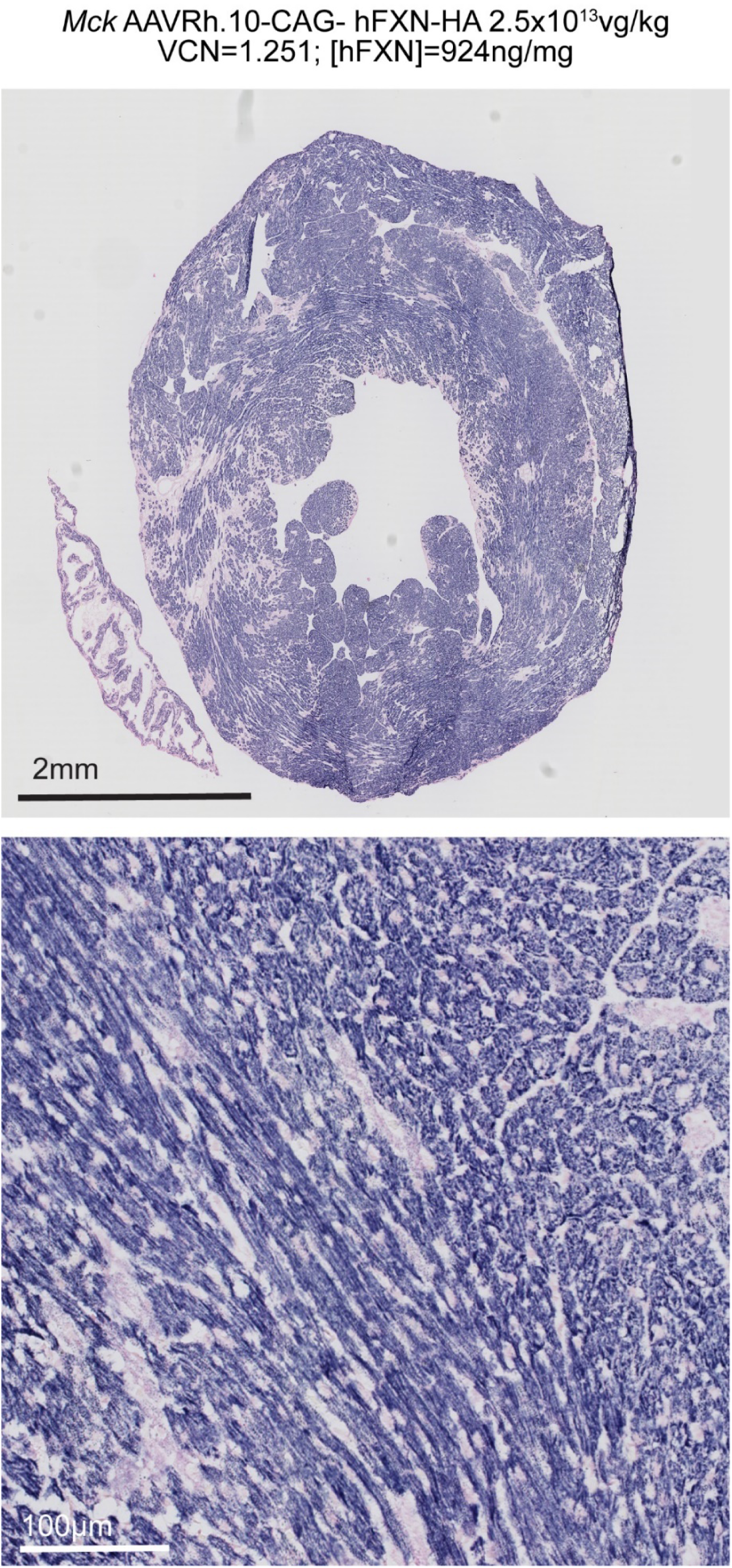
Rescue of SDH enzymatic activity in the heart of *Mck* mice treated with 2.5×10^13^ vg/kg of AAVRh.10-CAG-hFXN-HA vector. *In situ* histoenzymatic assay performed on heart tissue section from *Mck* mouse treated at 7 weeks of age and sacrificed at 15 weeks, and imaged at low and high magnifications.

**Table S1.** Statistical analysis of echocardiography measurements in *Mck* mice treated at 5 weeks of age with AAVRh.10-CAG-hFXN-HA vector at the two highest dose. One-way ANOVA analysis, Mixed-effects model (REML), no assumption of sphericity, α = 0.05

| Left ventricle shortening fraction |  |  |  |
| --- | --- | --- | --- |
| Fixed effect (type III) <0.0001 | F (DFn, DFd) F (1.135, 9.079) = 49.42 | Geisser-Greenhouse's epsilon = 0.3783 |  |
| Tukey's multiple comparisons test | Mean Diff. | 95.00% CI of diff. | Adjusted P Value |
| WT NaCl (n=7) vs. *Mck* 5x10<sup>13</sup> vg/kg (n=3) | 4.250 | 1.743 to 6.757 | **0.0044** |
| WT NaCl (n=7) vs. *Mck* 2.5x10<sup>13</sup> vg/kg (n=2) | 1.869 | 0.02753 to 3.711 | **0.0471** |
| WT NaCl (n=7) vs. *Mck* untreated (n=10) | 18.82 | 10.45 to 27.19 | **0.0016** |
| *Mck* 5x10<sup>13</sup> vg/kg (n=3) vs. *Mck* 2.5x10<sup>13</sup> vg/kg (n=2) | -2.381 | -4.643 to -0.1192 | **0.0406** |
| *Mck* 5x10<sup>13</sup> vg/kg (n=3) vs. *Mck* untreated (n=10) | 14.57 | 4.878 to 24.26 | **0.0123** |
| *Mck* 2.5x10<sup>13</sup> vg/kg (n=2) vs. *Mck* untreated (n=10) | 16.95 | 7.139 to 26.76 | **0.0074** |
| Cardiac output normalized to body weight |  |  |  |
| Fixed effect (type III) <0.0001 | F (DFn, DFd) F (1.590, 12.72) = 90.72 | Geisser-Greenhouse's epsilon = 0.5299 |  |
| Tukey's multiple comparisons test | Mean Diff. | 95.00% CI of diff. | Adjusted P Value |
| WT NaCl (n=7) vs. *Mck* 5x10<sup>13</sup> vg/kg (n=3) | 0.09299 | -0.4230 to 0.6090 | 0.9207 |
| WT NaCl (n=7) vs. *Mck* 2.5x10<sup>13</sup> vg/kg (n=2) | 0.2953 | -0.2019 to 0.7926 | 0.2669 |
| WT NaCl (n=7) vs. *Mck* untreated (n=10) | 2.727 | 1.868 to 3.586 | **0.0003** |
| *Mck* 5x10<sup>13</sup> vg/kg (n=3) vs. *Mck* 2.5x10<sup>13</sup> vg/kg (n=2) | 0.2024 | -0.08695 to 0.4917 | 0.1728 |
| *Mck* 5x10<sup>13</sup> vg/kg (n=3) vs. *Mck* untreated (n=10) | 2.634 | 1.374 to 3.894 | **0.0036** |
| *Mck* 2.5x10<sup>13</sup> vg/kg (n=2) vs. *Mck* untreated (n=10) | 2.432 | 1.454 to 3.410 | **0.0019** |
| Left ventricle mass normalized to body |  |  |  |
| Fixed effect (type III) = 0.0014 | F (DFn, DFd) F (1.015, 8.123) = 22.10 | Geisser-Greenhouse's epsilon = 0.3385 |  |
| Tukey's multiple comparisons test | Mean Diff. | 95.00% CI of diff. | Adjusted P Value |
| WT NaCl (n=7) vs. *Mck* 5x10<sup>13</sup>vg/kg (n=3) | 7.256e-005 | -2.800e-005 to 0.0001731 | 0.1575 |
| WT NaCl (n=7) vs. *Mck* 2.5x10<sup>13</sup>vg/kg (n=2) | 8.208e-005 | -2.791e-006 to 0.0001670 | 0.0571 |
| WT NaCl (n=7) vs. *Mck* untreated (n=10) | -0.001551 | -0.003006 to -9.562e-005 | 0.0395 |
| *Mck* 5x10<sup>13</sup>vg/kg (n=3) vs. *Mck* 2.5x10<sup>13</sup>vg/kg (n=2) | 9.524e-006 | -9.986e-005 to 0.0001189 | 0.9895 |
| *Mck* 5x10<sup>13</sup>vg/kg (n=3) vs. *Mck* untreated (n=10) | -0.001624 | -0.002941 to -0.0003064 | **0.0249** |
| *Mck* 2.5x10<sup>13</sup>vg/kg (n=2) vs. *Mck* untreated (n=10) | -0.001633 | -0.003034 to -0.0002318 | **0.0302** |
| Left ventricle end diastole diameter |  |  |  |
| Fixed effect (type III) = 0.0023 | F (DFn, DFd) F (1.074, 6.086) = 24.40 | Geisser-Greenhouse's epsilon = 0.3580 |  |
| Tukey's multiple comparisons test | Mean Diff. | 95.00% CI of diff. | Adjusted P Value |
| WT NaCl (n=7) vs. *Mck* 5x10<sup>13</sup>vg/kg (n=3) | 0.000 | -0.01115 to 0.01115 | >0.9999 |
| WT NaCl (n=7) vs. *Mck* 2.5x10<sup>13</sup>vg/kg (n=2) | 0.01476 | 0.005639 to 0.02388 | **0.0056** |
| WT NaCl (n=7) vs. *Mck* untreated (n=10) | -0.05041 | -0.09929 to -0.001542 | **0.0446** |
| *Mck* 5x10<sup>13</sup>vg/kg (n=3) vs. *Mck* 2.5x10<sup>13</sup>vg/kg (n=2) | 0.01476 | -0.0006284 to 0.03015 | 0.0589 |
| *Mck* 5x10<sup>13</sup>vg/kg (n=3) vs. *Mck* untreated (n=10) | -0.05041 | -0.09703 to -0.003798 | **0.0387** |
| *Mck* 2.5x10<sup>13</sup>vg/kg (n=2) vs. *Mck* untreated (n=10) | -0.06518 | -0.1239 to -0.006452 | **0.0355** |
| Left ventricle end systole diameter |  |  |  |
| Fixed effect (type III) = 0.0016 | F (DFn, DFd) F (1.010, 5.721) = 31.31 | Geisser-Greenhouse's epsilon = 0.3365 |  |
| Tukey's multiple comparisons test | Mean Diff. | 95.00% CI of diff. | Adjusted P Value |
| WT NaCl (n=7) vs. *Mck* 5x10<sup>13</sup>vg/kg (n=3) | -0.01839 | -0.02474 to -0.01205 | 0.0002 |
| WT NaCl (n=7) vs. *Mck* 2.5x10<sup>13</sup>vg/kg (n=2) | 0.001131 | -0.008187 to 0.01045 | 0.9729 |
| WT NaCl (n=7) vs. *Mck* untreated (n=10) | -0.1221 | -0.2032 to -0.04113 | 0.0097 |
| *Mck* 5x10<sup>13</sup>vg/kg (n=3) vs. *Mck* 2.5x10<sup>13</sup>vg/kg (n=2) | 0.01952 | 0.008714 to 0.03033 | 0.0032 |
| *Mck* 5x10<sup>13</sup>vg/kg (n=3) vs. *Mck* untreated (n=10) | -0.1037 | -0.1842 to -0.02330 | 0.0213 |
| *Mck* 2.5x10<sup>13</sup>vg/kg (n=2) vs. *Mck* untreated (n=10) | -0.1233 | -0.2138 to -0.03279 | 0.0176 |

**Table S2.** Statistical analysis of echocardiography measurements in *Mck* mice treated at 7 weeks of age with AAVRh.10-CAG-hFXN-HA vector or AAVRh.10-hFXN vector at 2.5×10^13^vg/kg. One-way ANOVA analysis, Mixed-effects model (REML), no assumption of sphericity, α = 0.05

| Left ventricle shortening fraction |  |  |  |
| --- | --- | --- | --- |
| Fixed effect (type III) = 0.0112 | F (DFn, DFd): F (0.7212, 3.366) = 27.54 | Geisser-Greenhouse's epsilon = 0.2404 |  |
| Tukey's multiple comparisons test | Mean Diff. | 95.00% CI of diff. | Adjusted P Value |
| *Mck* AAVRh.10-hFXN (n=8) vs. *Mck* AAVRh.10-CAG-hFXN-HA (n=6) | -1.992 | -9.261 to 5.278 | 0.6098 |
| *Mck* AAVRh.10-hFXN (n=8) vs. *Mck* untreated (n=10) | 14.00 | -125.7 to 153.7 | 0.4786 |
| *Mck* AAVRh.10-hFXN (n=8) vs. WT NaCl (n=8) | -9.949 | -24.29 to 4.392 | 0.1251 |
| *Mck* AAVRh.10-CAG-hFXN-HA (n=6) vs. *Mck* untreated (n=10) | 16.00 | -4.653 to 36.64 | 0.0960 |
| *Mck* AAVRh.10-CAG-hFXN-HA (n=6) vs. WT NaCl (n=8) | -7.957 | -14.23 to -1.683 | **0.0153** |
| *Mck* untreated (n=10) vs. WT NaCl (n=8) | -23.95 | -27.96 to -19.95 | **0.0003** |
| Cardiac output normalized to body weight |  |  |  |
| Fixed effect (type III) = 0.0112 | F (DFn, DFd): F (0.7212, 3.366) = 27.54 | Geisser-Greenhouse's epsilon = 0.2404 |  |
| Tukey's multiple comparisons test | Mean Diff. | 95.00% CI of diff. | Adjusted P Value |
| *Mck* AAVRh.10-hFXN (n=8) vs. *Mck* AAVRh.10-CAG-hFXN-HA (n=6) | -0.4267 | -1.736 to 0.8823 | 0.5016 |
| *Mck* AAVRh.10-hFXN (n=8) vs. *Mck* untreated (n=10) | 2.016 | -18.42 to 22.45 | 0.4847 |
| *Mck* AAVRh.10-hFXN (n=8) vs. WT NaCl (n=8) | -0.6183 | -2.992 to 1.756 | 0.6401 |
| *Mck* AAVRh.10-CAG-hFXN-HA (n=6) vs. *Mck* untreated (n=10) | 2.443 | 0.3170 to 4.569 | 0.0344 |
| *Mck* AAVRh.10-CAG-hFXN-HA (n=6) vs. WT NaCl (n=8) | -0.1916 | -1.029 to 0.6456 | 0.8813 |
| *Mck* untreated (n=10) vs. WT NaCl (n=8) | -2.635 | -3.164 to -2.105 | **0.0006** |
| Left ventricle mass normalized to body |  |  |  |
| Fixed effect (type III) = 0.0138 | F (DFn, DFd): F (0.7034, 3.282) = 24.91 | Geisser-Greenhouse's epsilon = 0.2345 |  |
| Tukey's multiple comparisons test | Mean Diff. | 95.00% CI of diff. | Adjusted P Value |
| *Mck* AAVRh.10-hFXN (n=8) vs. *Mck* AAVRh.10-CAG-hFXN-HA (n=6) | 0.0009792 | 4.837e-005 to 0.001910 | **0.0437** |
| *Mck* AAVRh.10-hFXN (n=8) vs. *Mck* untreated (n=10) | 0.001687 | -6.490e-006 to 0.003381 | **0.0505** |
| *Mck* AAVRh.10-hFXN (n=8) vs. WT NaCl (n=8) | -0.0004650 | -0.02119 to 0.02026 | 0.9513 |
| *Mck* AAVRh.10-CAG-hFXN-HA (n=6) vs. *Mck* untreated (n=10) | 0.0007082 | 0.0002797 to 0.001137 | **0.0033** |
| *Mck* AAVRh.10-CAG-hFXN-HA (n=6) vs. WT NaCl (n=8) | -0.001444 | -0.003638 to 0.0007495 | 0.1412 |
| *Mck* untreated (n=10) vs. WT NaCl (n=8) | -0.002152 | -0.003301 to -0.001004 | **0.0087** |
| Left ventricle end diastole diameter |  |  |  |
| Fixed effect (type III) = 0.0132 | F (DFn, DFd): F (0.7051, 5.170) = 15.30 | Geisser-Greenhouse's epsilon = 0.2350 |  |
| Tukey's multiple comparisons test | Mean Diff. | 95.00% CI of diff. | Adjusted P Value |
| *Mck* AAVRh.10-hFXN (n=8) vs. *Mck* AAVRh.10-CAG-hFXN-HA (n=6) | -0.001386 | -0.05710 to 0.05433 | 0.9992 |
| *Mck* AAVRh.10-hFXN (n=8) vs. *Mck* untreated (n=10) | -0.03596 | -0.9011 to 0.8292 | 0.8200 |
| *Mck* AAVRh.10-hFXN (n=8) vs. WT NaCl (n=8) | 0.02581 | -0.05129 to 0.1029 | 0.4848 |
| *Mck* AAVRh.10-CAG-hFXN-HA (n=6) vs. *Mck* untreated (n=10) | -0.03457 | -0.09014 to 0.02099 | 0.1603 |
| *Mck* AAVRh.10-CAG-hFXN-HA (n=6) vs. WT NaCl (n=8) | 0.02720 | 0.01198 to 0.04241 | **0.0020** |
| *Mck* untreated (n=10) vs. WT NaCl (n=8) | -0.06177 | -0.09496 to -0.02858 | **0.0088** |
| Left ventricle end systole diameter |  |  |  |
| Fixed effect (type III) = 0.0132 | F (DFn, DFd): F (0.7051, 5.170) = 15.30 | Geisser-Greenhouse's epsilon = 0.2350 |  |
| Tukey's multiple comparisons test | Mean Diff. | 95.00% CI of diff. | Adjusted P Value |
| *Mck* AAVRh.10-hFXN (n=8) vs. *Mck* AAVRh.10-CAG-hFXN-HA (n=6) | -0.001386 | -0.05710 to 0.05433 | 0.9992 |
| *Mck* AAVRh.10-hFXN (n=8) vs. *Mck* untreated (n=10) | -0.03596 | -0.9011 to 0.8292 | 0.8200 |
| *Mck* AAVRh.10-hFXN (n=8) vs. WT NaCl (n=8) | 0.02581 | -0.05129 to 0.1029 | 0.4848 |
| *Mck* AAVRh.10-CAG-hFXN-HA (n=6) vs. *Mck* untreated (n=10) | -0.03457 | -0.09014 to 0.02099 | 0.1603 |
| *Mck* AAVRh.10-CAG-hFXN-HA (n=6) vs. WT NaCl (n=8) | 0.02720 | 0.01198 to 0.04241 | **0.0020** |
| *Mck* untreated (n=10) vs. WT NaCl (n=8) | -0.06177 | -0.09496 to -0.02858 | **0.0088** |

**Table S3.**
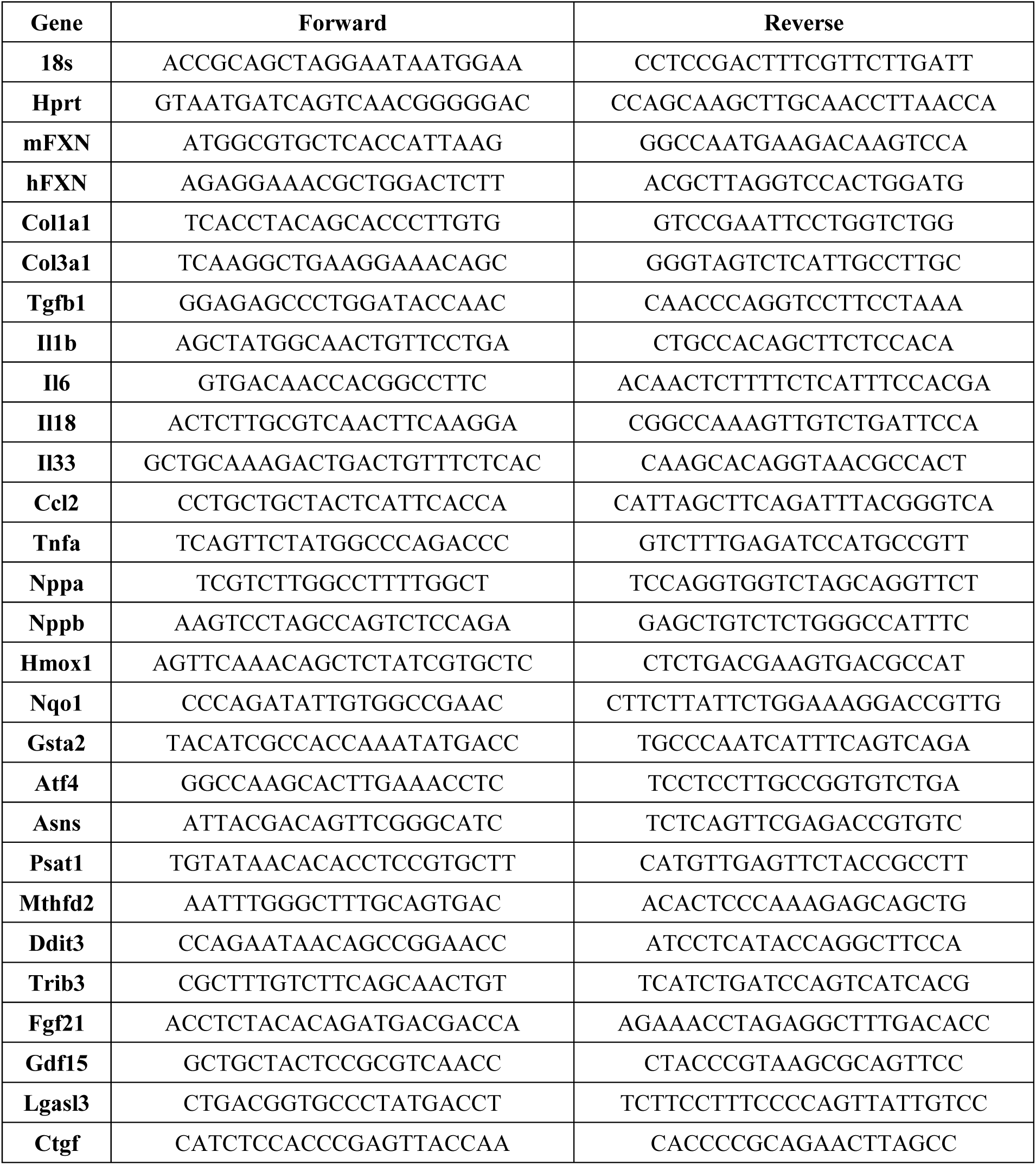
Primers sequences.

## SUPPLEMENTAL METHODS

### Adeno-associated viral vector constructs and production

The AAVRh.10-CAG-hFXN-HA and AAVRh.10-hFXN vectors are described in Figure S9A-B, and encode respectively the human frataxin cDNA, including the mitochondrial targeting sequence, and fused or not in C-terminal to the HA tag. These vectors were produced by triple transfection method in HEK293^3^, respectively at the Vector Core at the University Hospital of Nantes (France) and the Belfer Gene Therapy Core facility at Weill Cornell Medical College.

Their respective concentration was measured as 8.7×10^13^ and 1.96×10^14^ vector genomes per mL (vg/mL), by qPCR. Their purity was checked with a combination of the following assay: potential protein contamination by SDS-PAGE and Coomassie Blue Staining, endotoxin contamination with Endosafe®-PTSTM LA (< 0.5 EU/m), sterility by growth in liquid and agar based test media (no contamination detected at 48H), and qPCR quantification of residual DNA material from AAV REP or Adenovirus plasmids.

### Animal procedures

WT C57/B6J mice were supplied from Charles River laboratoires France. *Mck* mice were generated in 100% C57BL/6J background, with a conditional deletion of *Fxn* gene in cardiac and skeletal muscle (*Mck*-Cre-*Fxn*^L3/L-)^ and genotyped, as described previously ^4^. Housing animal facility was controlled for temperature and humidity, with a 12 hours light/dark cycle and free access to water and a standard rodent chow (D03, SAFE, Villemoisson-sur-Orge, France). All animal procedures and experiments were approved by the local ethical committee for Animal Care and Use (Com’Eth HP/2012/09/27 and 5344). They were performed in accordance with the Guide for the Care and Use of Laboratory Animals (National Institutes of Health). Five- and seven-weeks old animals were anesthetized with isoflurane prior to the retro-orbital injection of vector, in 100µL bolus. Control littermate or WT mice were injected with equivalent volume of NaCl 0.9% solution. Survival and comorbidities were evaluated daily and body weight, weekly. Echocardiography were performed with the Sonos 5500 system (Hewlett Packard) with a 15 MHz linear transducer (15L6) or with the Vevo 2100 system (Fujifilm Visualsonics) with a 25 to 55 MHz transducer, as described previously ^1^. Briefly, echocardiography was performed under isoflurane anesthesia (0.5-1.5%) and 0.8 mL/min O_2_, as to maintain the heart rhythm between 450 and 550 bpm. Historical data were used as reference for untreated *Mck* mice survival, body weight and cardiac function Following intraperitoneal injection of ketamine-xylazine and blood sampling, mice were perfused with cooled NaCl 0.9%. and the heart and liver were retrieved quickly.

Briefly, the apex of the heart was sampled in cryotubes and flash frozen in liquid nitrogen, for subsequent DNA and RNA extraction and analysis, as described previously ^1^. Two small pieces from the left ventricle’s anterior wall (less than 1mm3) were sampled in Karnovsky’s fixative and processed for transmitted electron microscopy (TEM) morphological analysis. A thin transversal section (around 2mm thick) was cut at the middle level of the heart for protein analysis. The middle to the basis of the heart was embedded in OCT and frozen in isopentane chilled with liquid nitrogen for histological analysis.

The liver medial left lobe was sampled and fixed in PBS 1x formaldehyde 10%, dehydrated and then embedded in paraffin for histological analysis. Two small pieces from the left lateral lobe (less than 2mm3) were sampled, fixed in Karnovsky’s fixative for TEM analysis. The remaining lobes - lateral left, medial right, lateral right and caudate lobes – were sampled in a cryotube and flash frozen in liquid nitrogen for subsequent DNA, RNA and protein analysis.

The spleen was fixed in PBX 1x Formaldehyde 4%, embedded in OCT and frozen in isopentane chilled with liquid nitrogen for histological analysis.

### Histochemistry

Frozen sections from the heart and the spleen, and liver paraffine sections, were stained with hematoxylin-eosin (H&E). In addition, heart tissue sections were stained with wheat germ agglutinin conjugated with Alexa 488nm (WGA), or stained with DAB-enhanced Perls labelling, or used to perform *in-situ* histoenzymatic activity assay for succinate dehydrogenase or cytochrome C oxidase, as previously described^1^. NADH dehydrogenase *in-situ* histoenzymatic activity was performed accordingly to Luna et al ^5^.

Immunofluorescent labelling, with or without SDH co-labelling, was performed as described previously ^1^, with the following antibodies and dilutions: frataxin (anti-FXN, 1/50, IGBMC, FXN935 and anti-HA, 1/100, Abcam, Ab9110), prohibitin (1/300, Ab28172), Sqstm1 (1/300, 2C11, H00008878-M01, Millipore) and acetyl-lysine (1/200, Ab21623, ABCAM).

For labelling of CD3, CD14 and CD45 positive cells, tissue sections were fixed in 4% PFA for 5min, permeabilized in PBS1x 0.3% Triton X-100 at RT for 10min and then blocked with PBS, 1% NGS, 5% BSA, 0.2% Tween (PBS-NBT) for 30min at RT. Subsequently, tissues sections were incubated O/N at 4°C with the rabbit antibody against CD3 (1/100, ab16669, ABCAM) or CD14 (1/100, ab183322, ABCAM,) or CD45 (1/100, ab10558, ABCAM) and afterward with alexa fluor-488nm conjugated goat anti-rabbit antibody (1/300, Molecular Probes).

Imaging of the heart, liver and spleen tissue sections were performed with the Hamamatsu NanoZoomer 2.0 slide scanner.

### Electron microscopy analysis

The samples were fixed with 2.5% glutaraldehyde and 2.5% paraformaldehyde in cacodylate buffer (0.1 M, pH 7.4), washed 30 minutes in cacodylate buffer, post-fixed with 1% osmium tetroxide in 0.1M cacodylate buffer for 1 hour at 4°C and dehydrated through graded alcohol (50, 70, 90, and 100%) and propylene oxide for 30 minutes each. Samples were oriented and embedded in Epon 812. Semithin sections were cut at 2µm with the Leica Ultracut UCT ultramicrotome and stained with 1% Toluidine blue and 1% sodium borate. Ultrathin sections were cut at 70nm and contrasted with uranyl acetate and lead citrate. Electron microscopy observation and image acquisition were performed at 70kv with the Morgagni 268D electron microscope (FEI Electron Optics, Eindhoven, and the Netherlands) and equipped with the Mega View III camera (Soft Imaging System). The bismuth sodium tartrate staining of ferritin was performed as described previously ^6, 7^.

### DNA, RNA and protein analysis

DNA, RNA and protein extraction and quantification, as well as cDNA synthesis, were performed as described previously ^1^.

Briefly, absolute quantification of vector biodistribution was performed in triplicate and the quantitative RT-PCR in duplicate, using the Quantitect Sybr® Green CR kit (Qiagen, 1037795) and the Light Cycler 480 II (Roche Biosciences). Primers were used at 0.4µM final concentration and their sequence are reported in Table S3. qPCR amplification program: 1 cycle, 15min 95°C; 50 cycles, 15sec 94°C-30sec 60°C-30s 72°C; 1 cycle for melting curve.

SimpleStep Human Frataxin ELISA Kit (ABCAM, ab176112) and SimpleStep Mouse Frataxin ELISA kit (ABCAM, ab199078) were used to quantify the heart and liver tissue concentration in human and mouse frataxin respectively, following manufacturer instructions. For Western Blot analysis, between 10 to 30µg of total protein extract were loaded and migrated in 14% or 4-12% SDS PAGE gels (Novex, Nupage 4-12% Bis-Tris Midi Gels 26 wells). Following transfer onto nitrocellulose membranes (Amersham Protran 0.45µm NC), Red Ponceau staining (Sigma Aldrich, P7170-1L), and blocking, the membranes were immunoblotted O/N at 4°C with the following antibodies and dilution: mouse anti-frataxin (1/10,000, 4F9, IGBMC), mouse anti-complex II 30KDa subunit SDHb (1/10,000, 21A11AE7, Novex), mouse anti-glyceraldehyde-3-phosphate dehydrogenase (1/30,000, MAB374, Millipore) and/or mouse anti-beta-tubulin (1/10,000, IGBMC). After incubation for 2 hours at 4°C with goat anti-mouse IgG H+L (1/10,000, 115-035-146, Jackson Immunoresearch), the signal was revealed using the SuperSignal West Dura Extended Duration Substrate (ThermoSienctific, 34075) and imaged with the Amersham Imager 600 (GE Healthcare).

### Statistical, correlation and regression analysis

Unless otherwise specified, data are reported as mean ± standard deviation (SD). Statistical analyses were carried-out using GraphPad Prism 8.3 (GraphPad Software, La Jolla California USA) and methods are described in the figure legends.

